# Assessing measurement uncertainty at laboratory network scale and its impact on diagnostic performance in the absence of a gold standard: application to ELISA tests for *Coxiella burnetii in ruminants*

**DOI:** 10.64898/2026.09.10.750550

**Authors:** Laureline Rivière, Marie-Laure Delignette Muller, Myriam Prigent, Thibaut Lurier, Elodie Rousset

**Affiliations:** Université Clermont Auvergne, INRAE, VetAgro Sup, UMR EPIA, Saint-Genès-Champanelle 63122, France; Université de Lyon, INRAE, VetAgro Sup, UMR EPIA, Marcy l’Etoile 69280, France; Universite Claude Bernard Lyon 1, LBBE, UMR 5558, CNRS, VetAgro Sup, Villeurbanne 69622, France; ANSES, Sophia Antipolis Laboratory, Animal Q fever Unit, Sophia Antipolis, France

**Keywords:** Batch, Bayesian, Q fever, Sensitivity, Specificity

## Abstract

Measurement uncertainty can affect the classifications of individual as positive or negative, and thus, the diagnostic performances of a test. Existing methods to assess measurement uncertainty and its impact on diagnostic performance are difficult to apply in the absence of a gold standard. We proposed a method applicable to any quantitative diagnostic test and in the absence of a gold standard, and applied it to ELISA tests for Q fever serology in ruminants. We assessed measurement uncertainty using a mixed-effects model on data from an inter-laboratory proficiency testing. Then, we estimated the sensitivities and specificities accounting for measurement uncertainty. To do so, we combined, for each individual of a sample representative of the population, the probability of being truly seropositive and the probability of being positive when retested in another laboratory. We also estimated the sensitivity and specificity of each laboratory and of each batch, taking into account their bias. While the sensitivities and specificities of the ELISA tests were slightly affected by the measurement uncertainty, they varied between laboratories and between batches. This highlights the importance of harmonising analytical practices across laboratories and calibrating batches. The extent to which measurement uncertainty impacts diagnostic performance depends not only on the values of the inter- and intra-laboratory standard deviations, but also on the position of the cut-off in the population’s measurand distribution. Therefore, in addition to the analytical performance of a test, the position of the cut-off relative to the distribution of test value is an essential parameter to consider.

## 1 Introduction

Most diagnostic tests use biological measurements to classify individuals as positive or negative, based on whether their measurement is above or below a cut-off. However, these measurements are subject to variability due to biological and analytical factors. As a result, the measured value may fluctuate around the cut-off, potentially resulting in changes to an individual classification (positive or negative) (Chai *et al*., 2017; Loh *et al*., 2024). This variability can affect the diagnostic performance of a test and thus undermine confidence in its results. This issue is particularly critical in the context of surveillance, where consistent and comparable diagnostic results across laboratories are essential for reliable epidemiological monitoring and public health decision-making. Therefore, understanding and quantifying measurement uncertainty, especially at the cut-off, is a prerequisite for harmonised diagnostic performances and robust surveillance outcomes.

Several serological ELISA tests for *C. burnetii* in ruminants are commercially available in Europe. Their diagnostic performances vary considerably (Lurier *et al*., 2021), which can affect the agreement between results from different laboratories and limit the comparability of surveillance data. In a previous study (Rivière et al., 2025), we estimated species-specific optimal cut-offs to harmonise these tests, by maximising the agreement between their results. Some of these new cut-offs are much higher or lower than the manufacturers’ cut-offs. This raises concerns about whether the analytical and diagnostic performances of the test can be maintained at these new cut-offs. In particular, the optimal cut-off for one of the tests in goats is low (6.6 versus 30 for the manufacturer cut-off), which may potentially affect the specificity of this test if the measurement uncertainty around this cut-off is high. Moreover, there is no gold standard for the serological diagnosis of *C. burnetii*. Therefore, a method applicable in the absence of a gold standard is needed to assess measurement uncertainty and its impact on diagnostic performance at the optimal cut-offs.

Several methods have been proposed for evaluating the impact of measurement uncertainty on diagnostic performance. A review of the literature (Smith *et al*., 2019) showed that a common approach involves simulations in which the following steps are repeated: (i) collecting measurand values considered as the “true values”, either from empirical data of individuals with known status, or from the measurand value distributions in the infected and non-infected populations, (ii) calculating the “measured values” by incorporating measurement uncertainty, and (iii) estimating the impact on diagnostic performance, such as the change in misclassification rate, or in sensitivity and specificity. Alternatively, Petersen *et al*. (2005) took measurement uncertainty into account to estimate the probability that a test value would be above the cut-off for each interval of possible values of the measurand. They then summed these probabilities, weighted by the proportion of non-infected individuals in each interval, to estimate the proportion of false positives. Another approach was proposed by Chatzimichail and Hatjimihail (2020), who developed formulas describing sensitivity and specificity as a function of measurement uncertainty. These formulas only apply if the measurand follows a Gaussian distribution with known parameters in both the infected and non-infected populations. All these methods require knowledge of the measurand distribution in both infected and non-infected populations. This can be challenging in the absence of a gold standard, particularly since it is difficult to obtain a sample of individuals representative of the target population with a known status. However, even in the absence of a gold standard, it is possible to calculate an individual’s probability of being infected based on their test results using the Bayes’ theorem applied to the output of a latent class model (which allows for the estimation of the prevalence of diseases and the sensitivity and specificity of diagnostic tests). These probabilities were used by Olsen *et al*. (2022) to estimate sensitivities and specificities at different cut-offs in order to estimate ROC curves in the absence of a gold standard. These probabilities, as well as the probabilities of being above or below the cut-off knowing the measurement uncertainty, could be used to estimate sensitivities and specificities including measurement uncertainty in the absence of a gold standard.

Regardless of the chosen method for evaluating the impact of measurement uncertainty, it is necessary to assess the measurement uncertainty beforehand. The importance of evaluating measurement uncertainty, particularly around the cut-off, is now also recognised in updated standards or guidelines. The revised French AFNOR standard NF U47-019 (2024) provides practical guidance for assessing measurement uncertainty at the cut-off for ELISA tests used in animal health and for reporting results close to the cut-off. This aligns with broader recommendations such as those in ISO/TS 20914:2019 (medical laboratory) (ISO, 2019) and the Guide to the Expression of Uncertainty in Measurement (GUM) (BIPM *et al*., 2008), which provide frameworks for identifying and quantifying sources of measurement variability in diagnostic settings. Measurement variability can occur at different levels and lead to measurement uncertainty (Dimech *et al*., 2006; Waugh, 2021). Intra-laboratory variability (also called intermediate precision or intra-laboratory reproducibility) can occur due to changes in operator, incubation time or temperature, as well as the variability between plates and within the same plate. Other contributors may include kit storage and shelf life, regular equipment maintenance, and instrument or pipette calibration, all of which are typically managed within ISO 17025 (ISO & IEC, 2017) compliant quality management systems. Inter-laboratory variability (also called inter-laboratory reproducibility) can be due to differences in instrument calibration, storage conditions, staff training or standardised procedures between laboratories. In addition, variability attributable to the test kit itself, including variation between batches (manufacturing runs), can contribute to measurement uncertainty in ELISA platforms, through differences in critical reagents, coating concentrations or buffer compositions, and through calibrator-related uncertainty (Suchanek & Robouch, 2009).

Depending on the level at which measurement uncertainty is being assessed, the relevant sources of variability must be considered. To assess measurement uncertainty at the network laboratory scale, measurements of the same material are required across different laboratories (to assess uncertainty due to inter-laboratory variability), as well as multiple measurements within laboratories (to assess uncertainty due to intra-laboratory variability). Inter-laboratory comparisons, where identical samples are tested by different laboratories, can be a way of obtaining the necessary data. While these schemes are typically used to evaluate the performance of participating laboratories, to validate analytical methods or to certify external reference materials, they could also be used to assess uncertainty due to inter- and intra-laboratory measurement variability by including several replicates of the same material per laboratory.

When evaluating its impact on diagnostic performance, it is important to assess the measurement uncertainty specifically around the cut-off, since samples with test values close to it are most likely to change result (from positive to negative or vice versa) upon retesting (WOAH, 2024). Therefore, the measurement uncertainty near the cut-off has a greater potential to affect diagnostic accuracy than the uncertainty elsewhere on the measurement scale. Furthermore, measurement uncertainty may depend on the underlying measurand value, as observed in previous inter-laboratory trials (Rousset & Dufour, 2019), highlighting the need to use samples with test values close to the cut-off when assessing measurement uncertainty.

The aim of this study is: (i) to assess the measurement uncertainty due to inter- and intra-laboratory variability around the optimal cut-offs of three Q fever ELISA tests, using data from an inter-laboratory proficiency testing, and (ii) to evaluate the impact of the measurement uncertainty by re-estimating the sensitivity and specificity of the tests, while accounting for measurement uncertainty.

## 2 Materials

### 2.1 Diagnostic tests

The three ELISA tests used to detect antibodies specific for *C. burnetii* in ruminant serum were the IDEXX Q Fever Ab Test, the PrioCHECK™ Ruminant Q Fever Ab Plate Kit, and the ID Screen® Q fever Indirect Multi-species. These tests detect IgG, using different types of conjugates (secondary antibodies or protein G). The antigens used in each test are derived from strains isolated from different species (tick, sheep or cow). The cut-offs used in this article were those that maximised agreement between the tests (Rivière *et al*., 2025) rather than the manufacturers’ cut-offs. The specifications of these tests are detailed in Table 1.

**Table 1.** ELISA tests used in the study.

| Name used in the study | Test 1 | Test 2 | Test 3 |
| --- | --- | --- | --- |
| Commercial name | IDEXX Q fever Ab test | PrioCHECK™ Ruminant Q Fever Ab Plate Kit | ID Screen® Q fever indirect multi-species |
| Manufacturer | IDEXX | Thermofisher Scientific | Innovative Diagnostics |
| Strain used for antigen production | Nine Mile reference strain, originally isolated from a *Dermacentor andersoni* tick | Isolated from an ewe | Isolated from a cow |
| Conjugate | Secondary antibodies binding to ruminant IgG | Protein G (binding to IgG of diverse mammalian species) | Protein G (binding to IgG of diverse mammalian species) |
| Cut-offs maximising the agreement, in optical density ratio (Rivière *et al.*, 2025) | Cattle: 44.9 %<br>Goat: 6.6 %<br>Sheep: 26.2 % | Cattle: 33.2 %<br>Goat: 18.9 %<br>Sheep: 55.0 % | Cattle: 84.3 %<br>Goat: 49.9 %<br>Sheep: 89.7 % |

The outputs of the tests are expressed in optical density ratio (ODR). According to the manufacturer’s instructions, they are calculated using formula (1) for tests 1 and 2, and formula (2) for test 3.

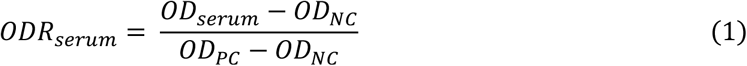

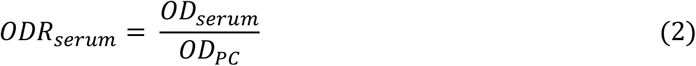

where *ODR*_*serum*_ is the optical density ratio of the tested serum, *OD*_*serum*_ is the optical density of the tested serum, and *OD*_*NC*_ and *OD*_*PC*_ are the optical density of the negative and positive controls included in the test, respectively.

### 2.2 Data

Two data sets were used in this study. The first one is from an inter-laboratory proficiency testing (ILPT) on Q fever ELISA tests. It was used to assess the measurement uncertainty. The second one, the Kiteval 4500 dataset, contains the ODRs measured with the three tests of a sample of cattle, goats and sheep that are representative of the population in a surveillance context. It was used to estimate the diagnostic performances.

#### Inter-laboratory proficiency testing (ILPT) data

To generate the ILPT data, sera were first prepared as follows: a pooled cattle serum from nine positive animals, a goat serum from a positive individual, and a sheep serum from a positive individual were each serially diluted (from 1/2 to 1/1024) in negative species-matched sera (foetal bovine serum, negative goat serum, and negative sheep serum, respectively). The French national Q fever laboratory then measured the ODR of each dilution using the three tests. For each species, one dilution was selected based on the proximity of its ODR values to the three optimal cut-offs recently proposed for the ELISA tests (Rivière *et al*., 2025). The three selected dilutions were prepared and distributed as lyophilised aliquots and were verified for homogeneity and stability by the organising laboratory in accordance with ISO 17043 (ISO, 2023) requirements, as described in Rousset & Prigent (2023). Hereafter, the terms “cattle serum”, “goat serum” and “sheep serum” will refer to the chosen diluted sera.

These three sera were included in the ILPT on ELISA methods for Q fever serology in ruminants organised by the French national Q fever laboratory in 2023 (Rousset & Prigent, 2023). Each participating laboratory received four samples of each of the three sera and tested them blindly using the ELISA test they routinely use. Several batches of ELISA tests were used during the ILPT, but each laboratory used a single batch for all analyses. The structure of the ILPT data is described in Figure 1.

**Figure 1.**
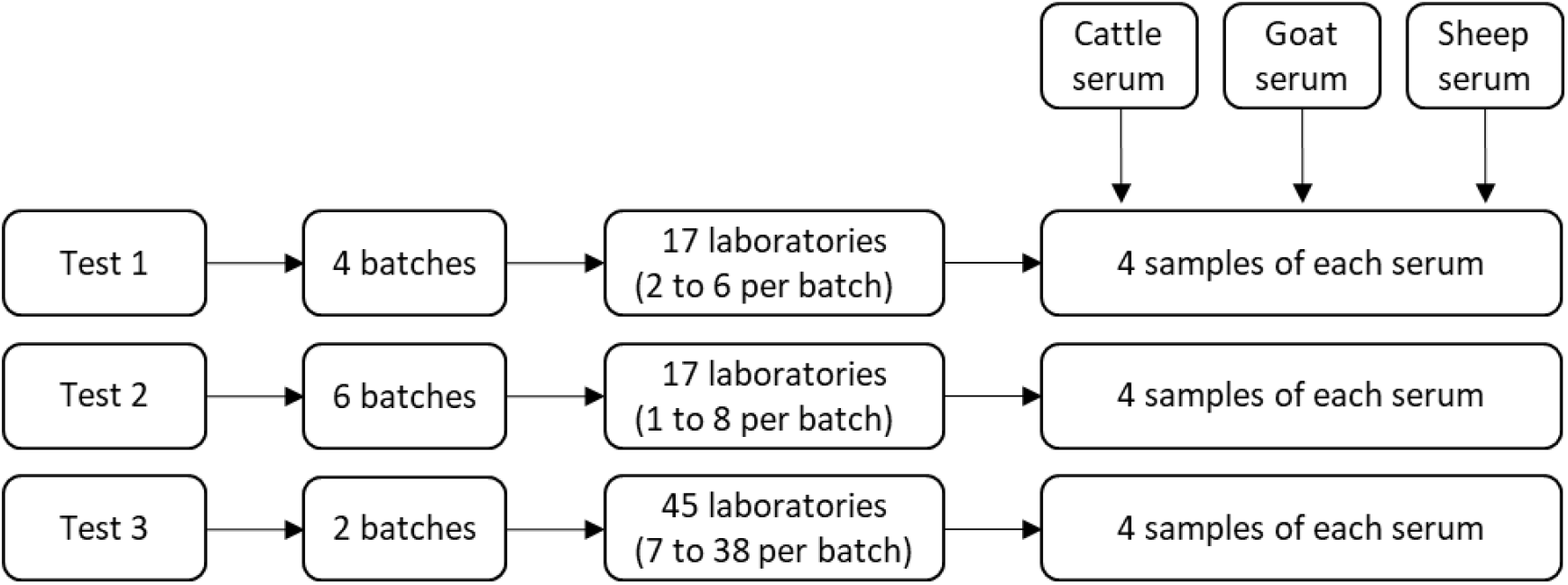
ILPT data structure.

#### Kiteval 4500 data

The Kiteval 4500 dataset used in this study is a previously analysed subsample from a national French serological survey on Q fever (Gache *et al*., 2017). Details on the sampling strategy and laboratory analyses can be found in Lurier *et al*. (2021). In brief, the dataset comprises sera from 1258 cattle, 1474 goats, and 1432 sheep, collected between 2012 and 2015 from 301 randomly selected herds across 10 French administrative departments. Each serum was analysed with the three ELISA tests by the French national Q fever laboratory. No new laboratory analyses were performed for the present work.

## 3 Methods

The statistical modelling was conducted in the following steps (see Figure 2): (i) The inter- and intralaboratory standard deviations (*σ*_*inter*_ and *σ*_*intra*_) were estimated using the ILPT data; (ii) These estimates were then used to calculate, for each individual in the Kiteval 4500 dataset, the probability of testing positive or negative when retested in another laboratory, referred to as the Retest Positive Probability (RPP) and Retest Negative Probability (RNP), respectively; (iii) Estimates from a latent class model applied to the Kiteval 4500 data were used to calculate the probability of each individual being truly seropositive, termed Posterior Positive Probability (PPP); (iv) The RPPs, RNPs and PPPs were then used to calculate the test sensitivities and specificities accounting for measurement uncertainty. These steps are more detailed in the following sections.

**Figure 2.**
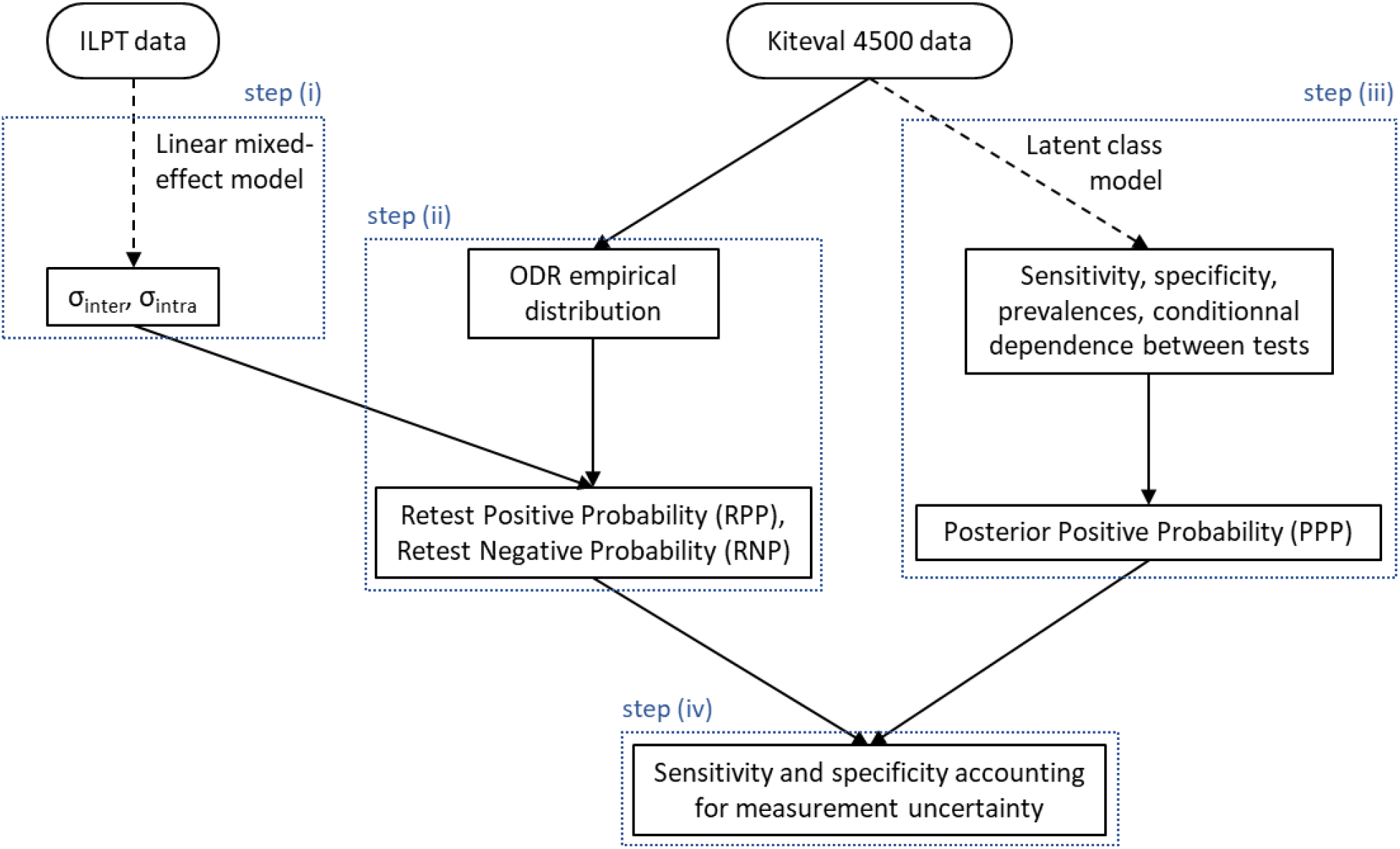
Modelling plan. The rounded squares represent the data sets and the squares represent the estimated parameters. The dashed arrows represent Bayesian inference and the solid arrows represent deterministic calculations.

These analyses were conducted independently for each test and species. Therefore, each step was performed nine times (once for each test-species combination).

All analyses were performed using `R` version 4.2.2. The `R` scripts used in this study can be found in the Supporting Information.

### 3.1 Variability assessment

#### Model description

For each test-serum combination, the ODR was modelled as a function of the laboratory and the batch using a linear mixed-effects model. The laboratory effect was described by a random effect (there are a large number of laboratories, 17 to 45 per test). On the other hand, the batch effect was described by a fixed effect, as recommended by van Leeuwen *et al*. (2022), since there are few different batches in the dataset (2 to 6 per test).

The ODR of sample *i* measured in laboratory *j* with batch *k* was modelled as follows:

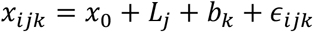

where *L*_*j*_ ~ Normal(0, *σ*_*inter*_) and *ϵ*_*ijk*_ ~ Normal(0, *σ*_*intra*_), with *x*_0_ representing the mean ODR of the serum, *L*_*j*_ the random effect of laboratory *j, b*_*k*_ the bias of batch *k* relative to the mean ODR of the serum measured across all batches, and *ϵ*_*ijk*_ the residual error. The inter-laboratory variability is described by the standard deviation of the random effects, *σ*_*inter*_, and the intra-laboratory variability by the residual standard deviation, *σ*_*intra*_.

#### Preliminary detection of outlier laboratories

As inter-laboratory proficiency testing schemes are designed to assess the proficiency of participating laboratories to produce results consistent with expected analytical performances, the presence of a limited number of outlier laboratories was expected. The leave-one-out method was used to determine if there were outlier laboratories. The following process was applied to each ILPT data subset corresponding to the nine test-serum combinations. The mixed-effects model described above was fitted to each data set obtained by removing one laboratory using a frequentist inference approach with the `lmer()` function from the `lme4` package (Bates *et al*., 2015) in `R`. At each iteration, the 95% confidence intervals of *σ*_*inter*_ and *σ*_*intra*_ were calculated using the `confint()` function with the default method. Then, the median of the lower and of the upper bounds of these confidence intervals was calculated. Laboratories for which, when removed, the point estimate of *σ*_*inter*_ or *σ*_*intra*_ was outside of these median bounds were excluded from all further analyses. These steps are illustrated in Figure A1 of the Supporting information for greater clarity.

After removing the outliers, we checked graphically that the residuals and random effects followed a Gaussian distribution centred on 0.

#### Inference (step i)

A Bayesian approach was used to estimate the parameters from the linear mixed-effects model on the data set from which the outlier laboratories were removed, in order to ensure methodological consistency with the subsequent estimation of the PPPs, which would be obtained via Bayesian inference (as explained below). This unified Bayesian framework facilitates the propagation of *σ*_*inter*_ and *σ*_*intra*_ uncertainty to the estimates of the sensitivity and specificity incorporating measurement uncertainty.

The priors used in the model are specified in Table 2, and the choice of the prior distributions is explained in Appendix A2 of the Supporting Information. The linear mixed-effects model was fitted to each subset of ILPT data corresponding to the ODR values observed for each of the 9 test-serum combinations. A Markov Chain Monte Carlo (MCMC) algorithm was run using three independent chains. For each chain, 110 000 iterations were produced. The first 10 000 were discarded as burn-in, and samples were then thinned by retaining every 15th iteration, resulting in 6 666 posterior samples per chain. The computations were performed using the `JAGS` software via the `R` package `rjags` (Plummer, 2008). The convergence was checked by displaying the MCMC chain traces and by computing the Gelman and Rubin’s statistics (Brooks & Gelman, 1998).

**Table 2.**
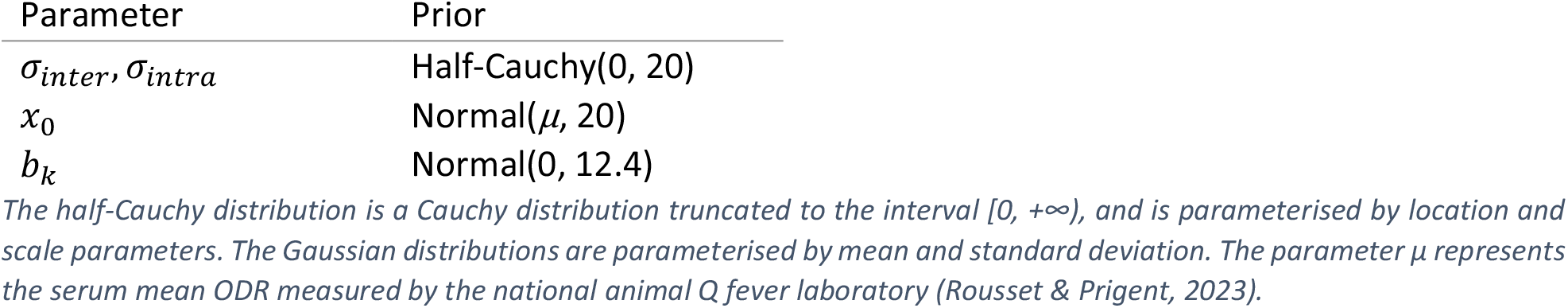
Prior distributions of unknown parameters.

### 3.2 Impact of the measurement variability on diagnostic performance

The Kiteval 4500 data were obtained from the reference laboratory for animal Q fever in France, which has the expertise in France for ruminants Q fever ELISA testing and follows all quality procedures standard: NF EN ISO IEC 17025 (ISO & IEC, 2017) and COFRAC’s LAB GTA 27 guide (Cofrac, 2020). Therefore, we considered the Kiteval 4500 data to be representative of the true ODR values of the individuals, and used them to estimate the sensitivities and specificities that do not account for measurement uncertainty. Sensitivities and specificities that included measurement uncertainty were estimated using the RPPs, RNPs and PPPs, and were then compared with the values obtained without measurement uncertainty. The method used is detailed in the following sections.

#### Probability of being negative or positive when retested (step ii)

To calculate the probability of a serum testing positive or negative when retested in another laboratory (RPP and RNP), the following hypotheses were made: the observed ODR in the Kiteval 4500 data (*x*_*obs*_) is considered to be the true serum ODR; the mean serum ODR in a laboratory follows a Gaussian distribution: *x*_*lab*_ ~ Normal(*x*_*obs*_, *σ*_*inter*_); the measured serum ODR follows a Gaussian distribution: *x* ~ Normal(*x*_*lab*_, *σ*_*intra*_).

The probability that a serum is negative when retested (RNP) i.e., that its new ODR *x* is below the cut-off *c* was calculated as follows:

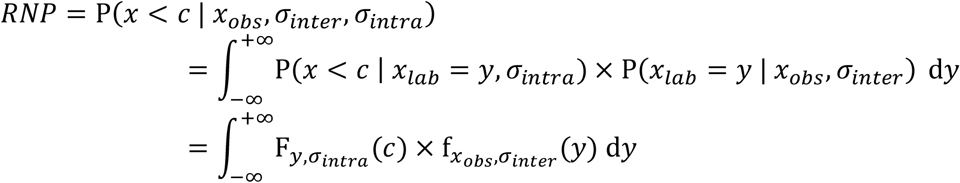

where 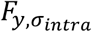 is the cumulative distribution function of a Gaussian distribution with mean *y* and standard deviation *σ*_*intra*_, and 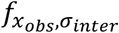 is the probability density function of a Gaussian distribution with mean *x*_*obs*_ and standard deviation *σ*_*inter*_.

The probability that a serum is positive when retested is:

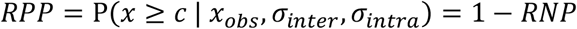

The integrals were calculated using the `hcubature()` function, with its default parameters, from the `cubature` (Narasimhan *et al*., 2025) `R` package.

#### Probability of being truly seropositive or seronegative (step iii)

The PPP (Posterior Positive Probability) of an individual can be expressed, according to Bayes theorem, as a function of the prevalence in its herd, of the sensitivities and specificities of the tests, and of the conditional dependence terms between the tests. These parameters were previously estimated on the Kiteval 4500 data using the latent class model described and assessed by Lurier *et al*. (2021).

Since individuals within the same herd and with the same combination of the three test results have the same PPP, the PPPs were calculated for each combination of herd and test results. The PPP formulas for each herd and test result combination can be found in Appendix A3 of the Supporting Information.

#### Sensitivity and specificity taking measurement uncertainty into account (step iv)

We used the formulas developed by Olsen *et al*. (2022) to estimate the sensitivities and specificities not accounting for measurement uncertainty, and we then adapted these formulas to estimate the sensitivities and specificities accounting for measurement uncertainty.

Olsen *et al*. (2022) proposed to estimate the sensitivity as the sum of the PPPs of the individuals tested positive (corresponding to the theoretical number of truly seropositive individuals tested positive) divided by the sum of the PPPs of all individuals (corresponding to the theoretical number of truly seropositive individuals). Similarly, the specificity can be calculated as the sum of the complements of the PPPs of the individuals tested negative (theoretical number of truly seronegative individual tested negative) divided by the sum of the complements of the PPPs of all individuals (theoretical number of truly seronegative individuals):

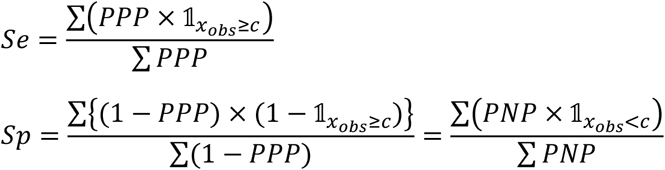

Where 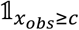 (respectively 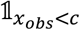) equals 1 for individuals tested positive (respectively negative) and 0 otherwise, and *PNP* = 1 − *PPP* is the probability of being truly seronegative (Posterior Negative Probability).

In order to take measurement uncertainty into account, we proposed, not to consider whether an individual result is positive or negative, but rather its probability of being positive or negative when retested in another laboratory. Therefore, we proposed to estimate the sensitivity (*Se*_*u*_) and the specificity (*Sp*_*u*_) accounting for measurement uncertainty as:

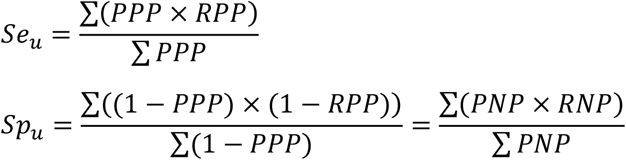

In these formulas, the numerator of *Se*_*u*_ corresponds to the theoretical number of truly seropositive individuals who would be positive when retested in another laboratory, and the numerator of *Sp*_*u*_ corresponds to the theoretical number of truly seronegative individuals who would be negative when retested.

Following the same principle, the number of false negatives and of false positives can be calculated as:

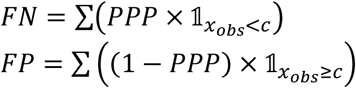

And the number of false negatives and of false positives taking into account the measurement uncertainty can be calculated as:

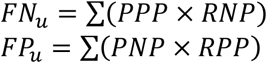

#### Sensitivity and specificity per laboratory

The sensitivity and specificity of each laboratory were also estimated. They do not take measurement uncertainty into account, but instead the laboratory effect. Each laboratory was considered to have a bias: its random effect. Therefore, an individual from Kiteval 4500 would test positive in laboratory *j* if its observed ODR (*x*_*obs*_) plus the laboratory bias (*L*_*j*_), was above the cut-off.

The formulas proposed by Olsen *et al*. (2022) were adapted by using the sum of the PPPs of the individuals from Kiteval 4500 who would test positive in laboratory *j* as the numerator of the sensitivity, and the sum of the PNPs of the individuals who would test negative in laboratory *j* as the numerator of the specificity. The sensitivity and specificity of laboratory *j* were calculated as follows:

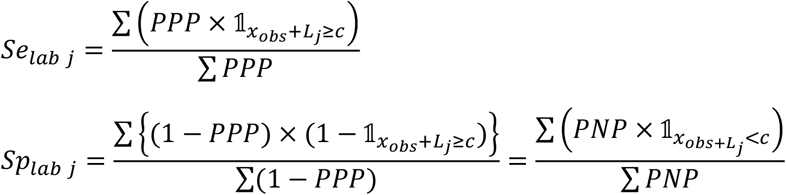

Where 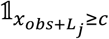 (respectively 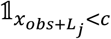) equals 1 for individuals for whom *x*_*obs*_ + *L*_*j*_ ≥ *c* (respectively *x*_*obs*_ + *L*_*j*_ < *c*) and 0 otherwise.

#### Sensitivity and specificity per batch

The sensitivities and specificities of each batch were calculated using the same method as for those of each laboratory, using the batch effects (*b*) instead of the laboratory effects. The sensitivity and specificity of batch *k* can be calculated as:

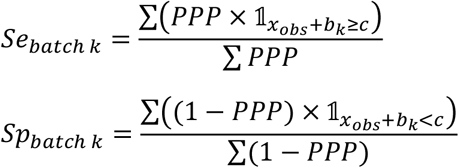

Where 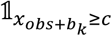 (respectively 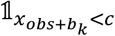) equals 1 for individuals for whom 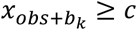 (respectively 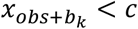) and 0 otherwise.

#### Point and interval estimation of parameters of interest

In order to estimate the 95% credibility intervals around the diagnostic performance, taking into account both the uncertainty surrounding the estimation of the parameters from the mixed model and that of the PPPs, the following steps were repeated 10 000 times: random draw of *σ*_*inter*_ and *σ*_*intra*_ and the laboratory effects *L*_*j*_ from their joint posterior distribution; random draw of the PPPs of each combination of test result and herd from their joint posterior distribution; and calculation of *Se, Sp, Se*_*u*_, *Sp*_*u*_, *FN, FP, FN*_*u*_, *FP*_*u*_, *Se*_*lab j*_, *Sp*_*lab j*_, *Se*_*batch k*_ and *Sp*_*batch k*_. The median, and 2.5% and 97.5% percentiles of the 10 000 estimates of each parameter were then calculated, respectively providing a point estimate and a 95% credibility interval.

## 4 Results

### 4.1 Outlier laboratories

Out of the 79 laboratories, three were identified as outliers. One laboratory that used test 1 and one that used test 3 were outliers for two species, and one laboratory that used test 2 was an outlier for the three species (Supporting Information – Figure A1.2).

### 4.2 Inter- and intra-laboratory standard deviations

Figure 3 shows the intra- and inter-laboratory standard deviations for each test-serum combination according to its mean ODR. For the three tests, higher values of ODR are associated with higher inter- and intra-laboratory standard deviations. The inter- and intra-laboratory standard deviations are both the lowest for the goat serum, which has the lowest ODR with the three tests.

**Figure 3.**
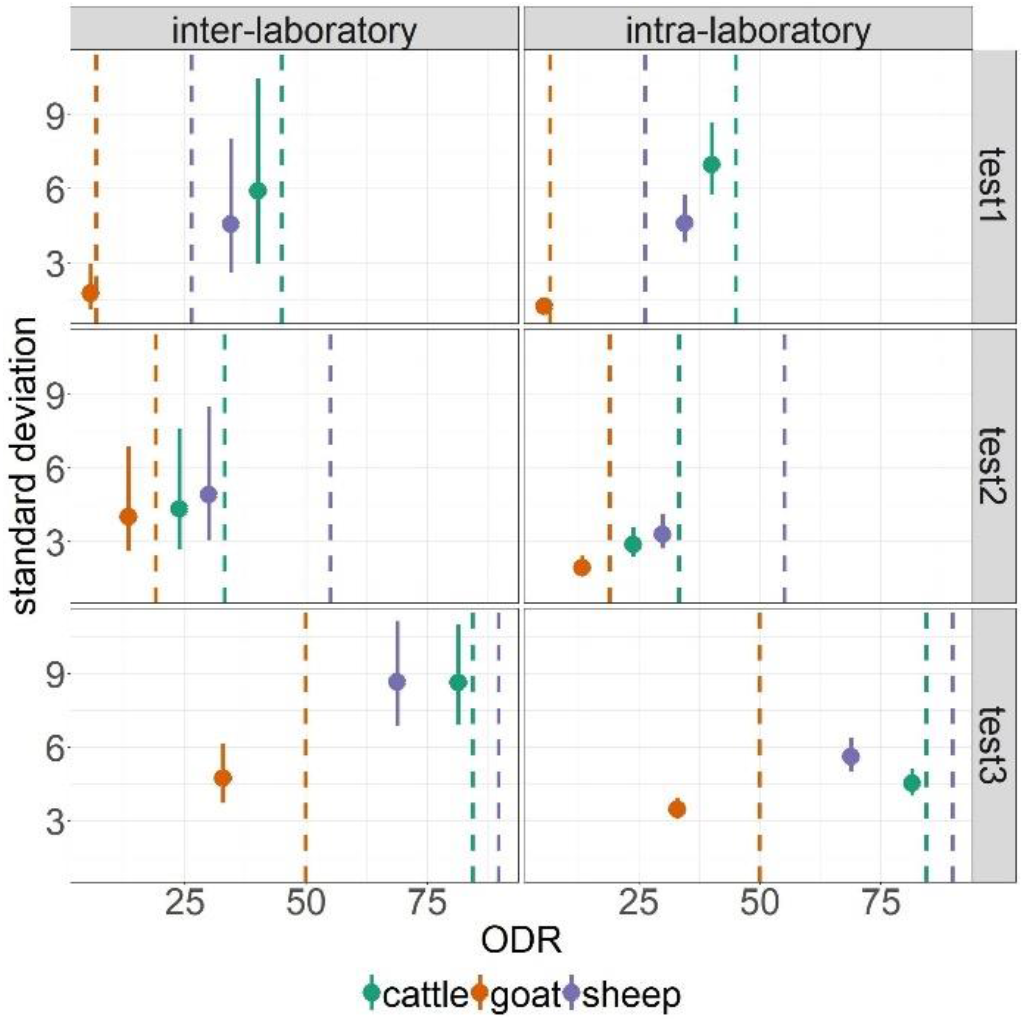
Inter- and intra-laboratory standard deviations of each serum according to their estimated mean ODR. The points represent their point estimate (median) and the solid lines their 95% credibility interval (2.5 and 97.5% quantiles). The dashed lines represent the cut-offs.

Overall, there is greater variability between laboratories than within laboratories. Indeed, the inter-laboratory standard deviations are higher than the intra-laboratory standard deviations, except for test 1 on the sheep serum, where they are similar, and for test 1 on the cattle serum, where the inter-laboratory standard deviation (5.8) is lower than the intra-laboratory standard deviation (7.0) (Figure 3).

### 4.3 Probability of changing result when retested in another laboratory

In order to see the proportion of individuals whose results are likely to change because of measurement uncertainty, Figure 4 shows the ODR distribution of the Kiteval 4500 individuals for the nine test-serum combinations, as well as the ODR value ranges for which there is a greater than 5% probability of obtaining different results when retested in another laboratory. The width of these ranges depends on the values of the inter- and intra-laboratory standard deviations. In particular, the range is the widest for test 3 in cattle and sheep, and the narrowest for test 1 in goats (Figure 4), corresponding to high and low values of intra- and inter-laboratory standard deviations, respectively (Figure 3). However, the proportion of individuals in these ranges is not necessarily high when the range is wide. Rather, it is high when the cut-off is close to the region of the ODR distribution with a high density of individuals, i.e., for tests and species with a low cut-off (Figure 4). Notably, this proportion is the highest in goats with test 1, which had the narrowest range.

**Figure 4.**
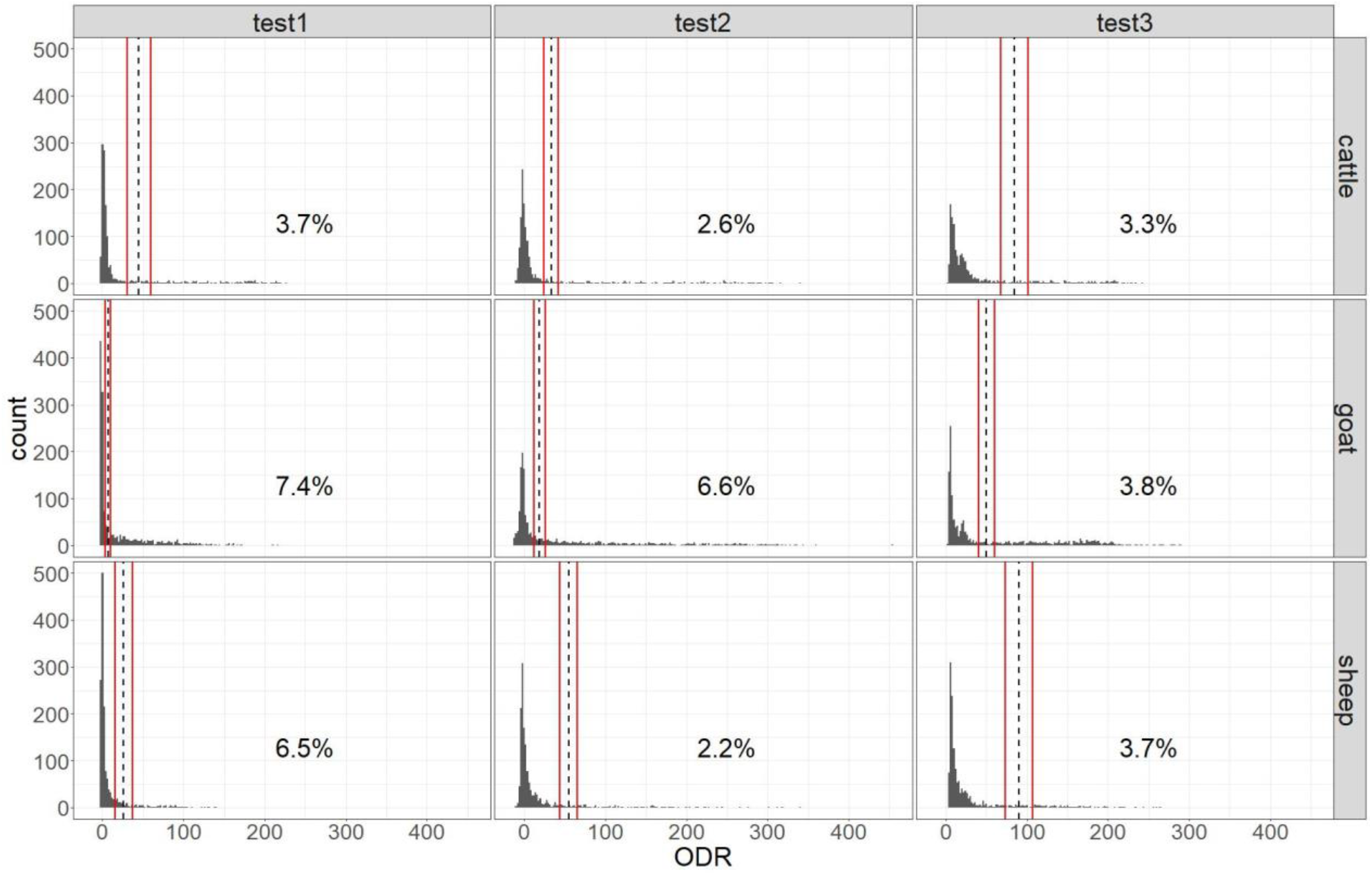
Proportion of Kiteval 4500 individuals with a probability of changing result greater than 5%. The solid red lines represent the range of ODRs for which the probability of changing result when retested in another laboratory is greater than 5%. The dashed lines represent the cut-offs. The histograms represent the ODR distribution in the Kiteval 4500 data

### 4.4 Impact of measurement uncertainty on diagnostic performance

The sensitivities and specificities estimated with and without accounting for measurement uncertainty are shown in Figure 5. The sensitivities and specificities accounting for measurement uncertainty (*Se*_*u*_ and *Sp*_*u*_) are generally lower than the unadjusted ones (*Se* and *Sp*). This difference is particularly noticeable for the specificity of test 1 in goats and sheep, and of test 2 in goats. However, *Se* and *Se*_*u*_ are very similar for test 1 in goats, test 2 in sheep and test 3 in goats, as are *Sp* and *Sp*_*u*_ for test 2 in sheep.

**Figure 5.**
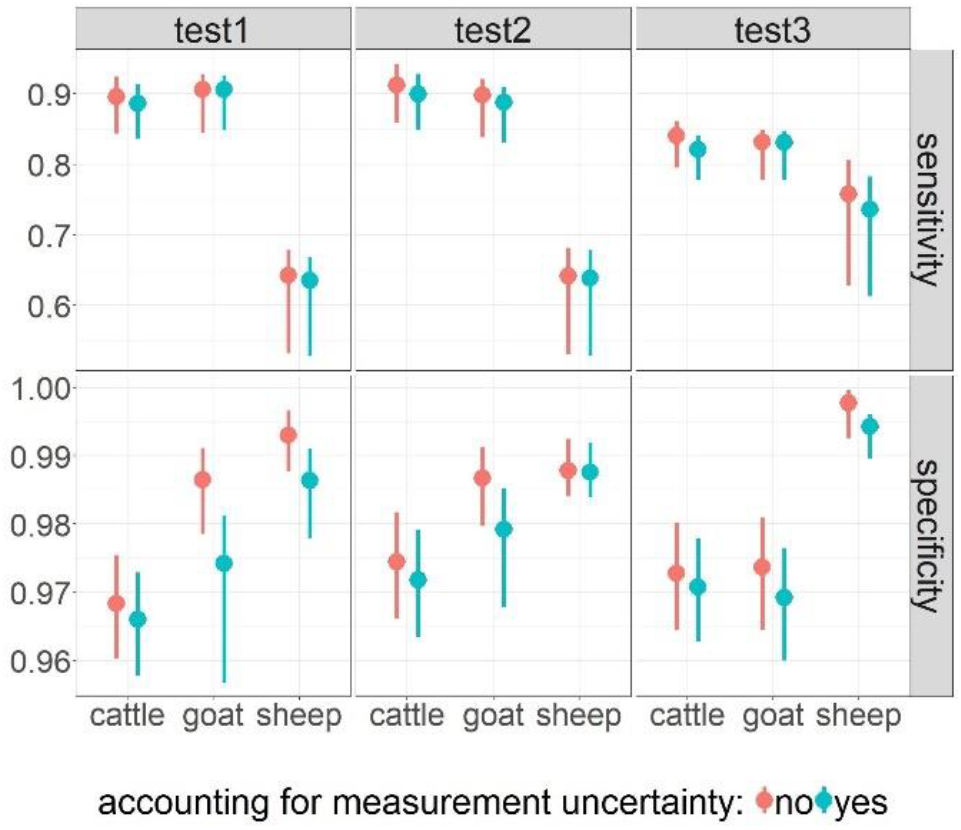
Sensitivities and specificities of the tests taking and not taking measurement uncertainty into account. The points represent their point estimate and the solid lines their 95% credibility interval.

When expressed as the number of misclassified individuals, accounting for measurement leads to an increase in the number of false negatives per 1000 individuals of 4.3 for test 2 in goats (from 41.9 to 46.2), and of 2.5 for test 3 in cattle and sheep (from 21.0 to 23.5, and from 30.4 to 33.1 respectively). For the six other test-species combinations, the increase is lower than 2 per 1000 individuals (Supporting Information – Table A4). The number of false positives per 1000 individuals increases by 7.2 per 1000 individuals for test 1 in goats (from 7.9 to 15.1), by 5.8 for test 1 in sheep (from 6.1 to 11.9), and by 4.4 for test 2 in goats (from 7.8 to 12.2). It increases by less than 3 per 1000 individuals for the other test-serum combinations.

Figure 6 illustrates the extent to which laboratory-specific sensitivities and specificities may depart from the unadjusted values, highlighting substantial heterogeneity between laboratories. When *Se*_*lab*._ is lower than *Se, Sp*_*lab*._ is higher than *Sp*, and vice versa. However, for a given laboratory, the magnitude of the difference between *Se*_*lab*._ and *Se* may differ from the magnitude of the difference between *Sp*_*lab*._ and *Sp*.

**Figure 6.**
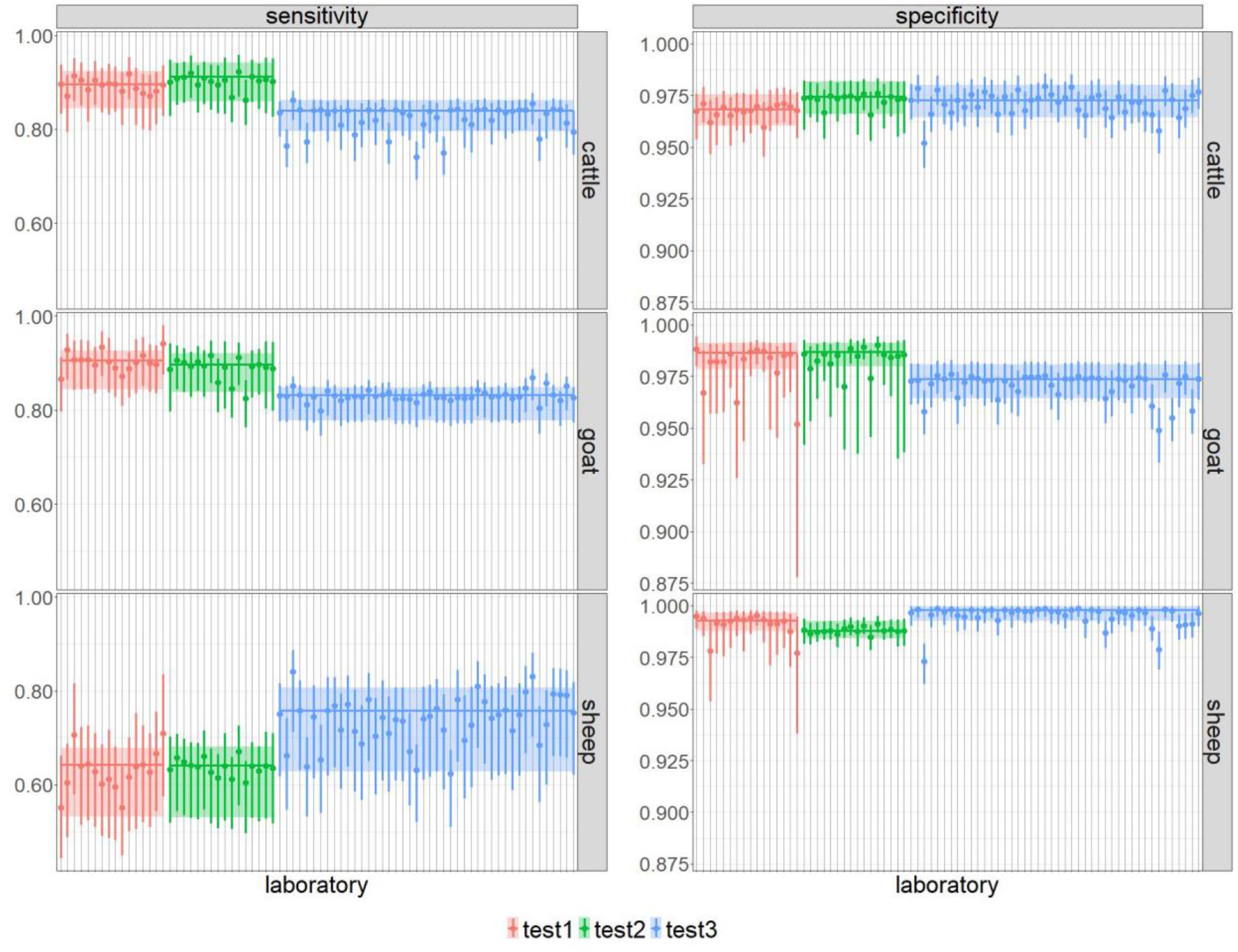
Sensitivity (Se_lab._) and specificity (Sp_lab._) of each of the 76 laboratories. Each point represents the point estimate of a laboratory and the vertical lines represent their 95% credibility intervals. The coloured horizontal lines and bands represent the point estimate and 95% credibility interval of the unadjusted sensitivity (Se) and specificity (Sp) of each test.

Some laboratory-adjusted sensitivities and specificities (*Se*_*lab*._ and *Sp*_*lab*._) differ greatly from their unadjusted counterparts (*Se* and *Sp*). For a given laboratory, they may differ greatly for one, two, or all three species.

The sensitivities, *Se*_*batch*._, and specificities, *Sp*_*batch*._, also vary between batches (Figure 7). As for the laboratories, when *Se*_*batch*._ is lower than *Se, Sp*_*batch*._ is higher than *Sp*, and vice versa. In particular, the diagnostic performances of the batches of test 1 in sheep are very different from the unadjusted ones. It is the case for the sensitivity of the four batches (two are much higher than *Se* and two much lower), as well as for the specificity of two of the four batches which are both much lower than *Sp*. The specificity of the other two batches is only slightly higher than *Sp*, given that *Sp* is close to 1.

**Figure 7.**
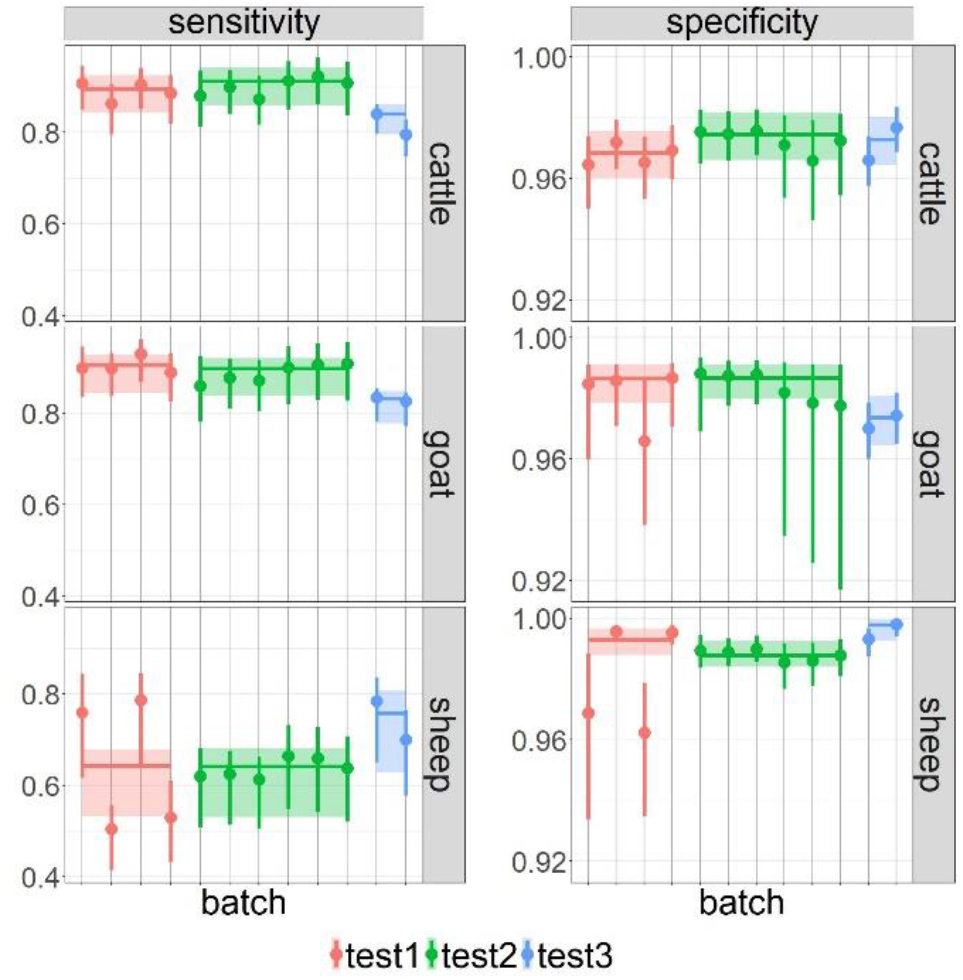
Sensitivity (Se_batch._) and specificity (Sp_batch._) of each batch. Each point represents the point estimate of a batch and the vertical lines represent their 95% credibility intervals. The coloured horizontal lines and bands represent the point estimate and 95% credibility interval of the unadjusted sensitivity (Se) and specificity (Sp) of each test.

## 5 Discussion

### 5.1 Main results

We have proposed a generic method for assessing the impact of measurement uncertainty on diagnostic performance. This method can be applied to any quantitative diagnostic test and in the absence of a gold standard. This approach is original because it integrates, within a single probabilistic and Bayesian framework, the main sources of variability: inter-laboratory, intra-laboratory, and batch-to-batch effects. Such a comprehensive treatment is rare in the literature on diagnostic test evaluation. It also aligns with recent standards that require explicit consideration of measurement uncertainty, particularly around diagnostic cut-offs (AFNOR, 2024; ISO, 2019). By enabling direct estimation of performance loss due to “real-life” uncertainties, this approach is a step towards more responsible and rigorous use of diagnostic data, particularly in surveillance and public health. The synergy between biostatistics (methodological innovation) and reference laboratory activities (practical implementation, network structuring, quality assurance) demonstrates the value of collaborative work between fundamental and applied research.

In this study, we applied this method to evaluate three commercial ELISA tests used for Q fever serology in ruminants. At the laboratory network scale, the diagnostic performances of the ELISA tests were slightly affected by their measurement uncertainty (Figure 5). However, they varied between laboratories (Figure 6) and between batches (Figure 7), highlighting the importance of harmonising analytical practices across laboratories and of calibrating the different batches of a test to ensure better data comparability.

When considering measurement uncertainty, it is essential to assess the variability of the test results around the cut-off, but also to quantify its impact on sensitivity and specificity. Indeed, the impact of measurement uncertainty on diagnostic performance depends, not only on the value of the standard deviations, but also on the proportion of individuals close to the cut-off. For example, compared to the other test-species combinations, test 1 in goats had the lowest intra- and inter-laboratory standard deviations (Figure 3), but it showed the largest decrease in specificity when measurement uncertainty was taken into account (Figure 5). Indeed, the cut-off of test 1 in goats is low and close to an area of the ODR distribution with a high density of individuals (Figure 4). Therefore, a large number of individuals with a “true” ODR below the cut-off may have a measured ODR above the cut-off because of the measurement uncertainty, which explains the decrease in specificity. This finding illustrates that measurement uncertainty has the greatest effect when the cut-off is set in an area with a high density of individuals. In such cases, even minor variations in measurement can lead to a substantial number of results shifting from one side of the cut-off to the other. Consequently, particular attention should be paid to the distribution of the measurand when selecting or adjusting cut-offs in routine practice.

### 5.2 Relation between ODR values and measurement uncertainty

There seems to be a positive correlation between the ODR value and the intra- and inter-laboratory standard deviations (Figure 3). It shows that it is important to assess the measurement uncertainty on sera close to the cut-off values, since individuals with a measurement close to the cut-off are more likely to change results. However, the ODR values of the sera used in our study to estimate intra- and inter-laboratory variabilities were not always as close to the cut-offs as expected, particularly for test 2 in sheep and test 3 in goats and sheep (Figure 3). The ODRs of these sera are below the cut-offs (respective ODRs of 29.9, 32.8 and 68.7, for cut-off values of 55.0, 49.9 and 89.7). Therefore, we may have underestimated the inter- and intra-laboratory variability around these cut-offs, and thus the probability of changing result when retested. However, this should have a limited impact on the estimation of the sensitivities and specificities accounting for measurement uncertainty, because these cut-offs are in an area of the ODR distributions with a low density of individuals (Figure 4), so only a small number of individuals is likely to change result.

Considering one test and one serum, we used the same standard deviations to calculate the probability of changing result of each individual, regardless of its ODR value. Thus, we probably overestimated the probability of changing result of the individuals with an ODR well below the cut-off and underestimated it for those with an ODR well above the cut-off. Our estimations would have been more accurate if we had modelled the inter- and intra-laboratory standard deviations as a function of the ODR, but this would have required many more sera to be included in the ILPT. However, as the individuals with an ODR close to the cut-off are the most likely to change result, it is more important to have an estimation of the standard deviations close to the cut-off.

### 5.3 Model hypotheses and limitations

The intra-laboratory standard deviation was modelled as constant across laboratories for one test and one species, while the width of the range of the values measured by a laboratory can differ greatly between laboratories (Supporting Information – Figure A5), indicating a potential variation in the intra-laboratory variability. However, estimating different standard deviations would have been difficult as there were only four measurements of each serum per laboratory.

To avoid biased estimates of the inter- and intra-laboratory standard deviations, we included the batch as a fixed effect in the mixed-effects model, as the batch may impact the measured ODR (Rousset *et al*., 2023). It would have been interesting to estimate an inter-batch standard deviation by modelling the batch effect as a random effect. However, only a small number of batches were used in the ILPT (2 to 6 per test) and inter-batch variability may change from one year to the next depending on the new batches commercialised. So, in any case, the estimated variability would not have been generalisable.

We calculated the PPPs using the qualitative results (positive or negative) of the individuals and not their ODR values. Therefore, an individual with ODRs close to the cut-offs, and thus more likely to change result, has the same PPP as an individual with ODRs far from the cut-offs, whereas its seropositivity status is probably less certain. This may lead to an overestimation of the impact of the measurement uncertainty on the sensitivity and specificity. However, estimating the probability of seropositivity while accounting for individuals ODR values would require knowledge of the ROC curves of the three tests, which are not currently available.

### 5.4 Impact of measurement variability

The sensitivities and specificities accounting for measurement uncertainty were almost all lower than the ones not accounting for it (Figure 5). The species and tests with the highest density of individuals around the cut-off were the most affected (Figures 4 and 5). This is coherent with the fact that accounting for measurement uncertainty adds a source of potential misclassification. However, the sensitivity accounting for measurement uncertainty of test 1 in goats is slightly higher. This can probably be explained by the fact that the cut-off of test 1 in goats is just above an area of the ODR distribution with a high density of individuals, so there are more truly seropositive individuals initially tested negative that become positive due to measurement uncertainty than truly seropositive individuals initially positive that become negative. This slight increase in sensitivity is also associated with an important decrease of specificity which is consistent with a high proportion of truly seronegative individuals that become positive because of measurement uncertainty.

The magnitude of the difference between *Se*_*lab j*_ and *Se*, or *Sp*_*lab j*_ and *Sp* may vary between species for a given laboratory (Figures 6), as well as the difference between *Se*_*batch k*_ and *Se*, or *Sp*_*batch k*_ and *Sp* for one batch (Figures 7). This magnitude depends on the proportion of individuals which may change result because of the laboratory or batch effect. This proportion depends on the size of the laboratory or batch effect, and on the density of individuals around the cut-off. These two parameters are affected by the cut-off value, which varies between species. Indeed, the inter-laboratory variability increases with the cut-off value, which may lead to higher laboratory effects, and batch effects appear to be generally higher for sera with a higher ODR (Supporting Information – Figure A5).

The sensitivities and specificities may vary greatly between laboratories, but also between batches. To our knowledge, the demonstration of such batch-related bias on sensitivity and specificity has not previously been documented in the literature on Q fever ELISAs. Notably, health authorities do not currently require formal validation or systematic batch-by-batch quality control for ELISA kits used in the diagnosis of Q fever in ruminants. Q fever remains classified as a neglected zoonosis, and the lack of mandatory standards for test validation or routine batch control represents a significant regulatory gap. Our findings highlight the necessity for precise tracking and documentation of the batch used for each test result, not only at the laboratory level but also for surveillance networks, and call for manufacturers to perform and communicate batch-to-batch evaluations.

### 5.5 Practical implications for ILPT planning

Our method can be a way to improve guidelines for the assessment of laboratory analytical performance. For semi-quantitative methods, such as ELISA tests, ILPTs are primarily designed to assess the capacity of laboratories to correctly classify sera. In the absence of a gold standard, reference laboratories generally include strongly positive or negative sera that are not representative of all the sera routinely analysed by laboratories. By estimating the measurement uncertainty around the cut-offs using ILPT data, and then the sensitivities and specificities in each laboratory using a sample representative of the target conditions and population, we can assess laboratory diagnostic performances that are closer to real conditions. In order to apply this method to a test, an ILPT must be designed to assess the measurement uncertainty around the cut-off of the test by including several samples of a serum close to the cut-off. The reference laboratory must also have access to quantitative test results from a sample representative of the population, and an estimation of the true sensitivity and specificity of the test (which can be obtained through a latent class model in the absence of a gold standard).

Moreover, when analysing ILPT data to assess the analytical performance of laboratories, it is important to take the batch into account. Otherwise, if a laboratory’s analytical performance is poor, it will be impossible to determine whether this is due to the batch or the laboratory’s analytical practices.

## 6 Conclusion

This article describes a new method to assess the impact of measurement uncertainty on diagnostic performance applicable in the absence of a gold standard. This method was applied to ELISA tests for the serological diagnosis of Q fever on ruminants. Although the sensitivity and specificity of these tests are only slightly affected at the laboratory network scale, they may vary greatly between laboratories and between batches. Moreover, it shows that, in addition to the inter- and intra-laboratory standard deviations, the position of the cut-off relative to the distribution of the biological marker in the population has a significant impact on the diagnostic performances. Therefore, it seems important to take this parameter into account when assessing analytical performance.

## Acknowledgements

The authors would like to thank the French platform for epidemiological surveillance in animal health, the farmers who participated in the Kiteval4500 project, the veterinarians who collected the samples, the Departmental Veterinary Laboratories that performed the analyses, and the Animal Health Farmers’ Organisations (GDS France) that coordinated the sample collection. We also thank the laboratories that participated to the ILPT, and Stéphane Groncin (ANSES) for information systems support and for aggregating and pre-curating the ILPT data.

This study was funded by VetAgro Sup, the French Agency for Food, Environmental and Occupational Health & Safety (ANSES).

## Data availability statement

The ILPT data are available in the supplementary material of this article.

The Kiteval 4500 data are available from the authors with the permission of GDS France.

## References

AFNOR. (2024). NF U47-019. Méthodes d’analyse en santé animale—Exigences et recommandations pour la validation, l’adoption et la mise en œuvre des techniques ELISA. Association Française de Normalisation. https://www.boutique.afnor.org/fr-fr/norme/nf-u47019/methodes-danalyse-en-sante-animale-exigences-et-recommandations-pour-la-val/fa205350/418514

Bates, D., Mächler, M., Bolker, B., & Walker, S. (2015). Fitting Linear Mixed-Effects Models Using lme4. Journal of Statistical Software, 67, 1–48. 10.18637/jss.v067.i01

BIPM, IEC, IFCC, ILAC, ISO, IUPAC, IUPAP, & OIML. (2008). Evaluation of measurement data—Guide to the expression of uncertainty in measurement. Joint Committee for Guides in Metrology, JCGM 100:2008. 10.59161/JCGM100-2008E

Brooks, S. P., & Gelman, A. (1998). General Methods for Monitoring Convergence of Iterative Simulations. Journal of Computational and Graphical Statistics, 7(4), 434–455. 10.1080/10618600.1998.10474787

Chai, J. H., Ma, S., Heng, D., Yoong, J., Lim, W.-Y., Toh, S.-A., & Loh, T. P. (2017). Impact of analytical and biological variations on classification of diabetes using fasting plasma glucose, oral glucose tolerance test and HbA1c. Scientific Reports, 7(1), 13721. 10.1038/s41598-017-14172-8

Chatzimichail, T., & Hatjimihail, A. T. (2020). A Software Tool for Exploring the Relation between Diagnostic Accuracy and Measurement Uncertainty. Diagnostics, 10(9), Article 9. 10.3390/diagnostics10090610

Cofrac. (2020). LAB GTA 27: Essais en immuno-sérologie animale (Rév. 03).). Comité Français d’Accréditation. https://tools.cofrac.fr/documentation/LAB-GTA-27

Dimech, W., Francis, B., Kox, J., & Roberts, G. (2006). Calculating Uncertainty of Measurement for Serology Assays by Use of Precision and Bias. Clinical Chemistry, 52(3), 526–529. 10.1373/clinchem.2005.056689

ISO. (2019). ISO/TS 20914:2019. Medical Laboratories—Practical Guidance for the Estimation of Measurement Uncertainty. International Organization for Standardization. https://www.iso.org/standard/69445.html

ISO. (2023). SO/IEC 17043:2023. Conformity assessment—General requirements for the competence of proficiency testing providers. https://www.iso.org/standard/80864.html

ISO, & IEC. (2017). ISO/IEC 17025. General requirements for the competence of testing and calibration laboratories. https://www.iso.org/fr/standard/66912.html

Loh, T. P., Markus, C., & Lim, C. Y. (2024). Impact of analytical imprecision and bias on patient classification. American Journal of Clinical Pathology, 161(1), 4–8. 10.1093/ajcp/aqad115

Lurier, T., Rousset, E., Gasqui, P., Sala, C., Claustre, C., Abrial, D., Dufour, P., de Crémoux, R., Gache, K., Delignette-Muller, M.-L., Ayral, F., & Jourdain, E. (2021). Evaluation using latent class models of the diagnostic performances of three ELISA tests commercialized for the serological diagnosis of Coxiella burnetii infection in domestic ruminants. Veterinary Research, 52(1), 56. 10.1186/s13567-021-00926-w

Narasimhan, B., Koller, M., Eddelbuettel, D., Johnson, S. G., Hahn, T., Bouvier, A., Kiêu, K., & Gaure, S. (2025). cubature: Adaptive Multivariate Integration over Hypercubes (Version 2.1.4) [Computer software]. https://cran.r-project.org/web/packages/cubature/index.html

Olsen, A., Nielsen, H. V., Alban, L., Houe, H., Jensen, T. B., & Denwood, M. (2022). Determination of an optimal ELISA cut-off for the diagnosis of Toxoplasma gondii infection in pigs using Bayesian latent class modelling of data from multiple diagnostic tests. Preventive Veterinary Medicine, 201, 105606. 10.1016/j.prevetmed.2022.105606

Petersen, P. H., Jørgensen Lone G. M., Brandslund,Ivan, De Fine Olivarius Niels, & and Stahl, M. (2005). Consequences Of Bias and Imprecision in Measurements of Glucose and Hba1c for the Diagnosis and Prognosis of Diabetes Mellitus. Scandinavian Journal of Clinical and Laboratory Investigation, 65(sup240), 51–60. 10.1080/00365510500236135

Plummer, M. (2008). rjags: Bayesian Graphical Models using MCMC [Data set]. The R Foundation. 10.32614/cran.package.rjags

Rivière, L., Rousset, E., Jourdain, E., Delignette-Muller, M.-L., & Lurier, T. (2025). Harmonisation of the diagnostic performances of serological ELISA tests for C. burnetii in ruminants in the absence of a gold standard: Optimal cut-offs and performances reassessment. Preventive Veterinary Medicine, 239, 106509. 10.1016/j.prevetmed.2025.106509

Rousset, E., Couesnon, A., Prigent, M., Jourdain, E., & Lurier, T. (2023). Analytical calibration of batches of ELISA kits used for the indirect diagnosis of Q fever in ruminants: Work in progress [ISWAVLD 2023]. ISWAVLD 2023. https://hal.inrae.fr/hal-04141750

Rousset, E., & Dufour, P. (2019). Inter-laboratory proficiency testing programme report (FQELSE19): Q fever serology with serum by ELISA. (No. V00; p. 50 pages). ANSES - Laboratoire de Sophia Antipolis, Unité fièvre Q animale (UFQa). https://hal.science/hal-05349536

Rousset, E., & Prigent, M. (2023). Final Inter-Laboratory Proficiency Testing Program Report (FQELSE23): Q fever serology on serum by ELISA (p. 43 pages) [Report]. ANSES. https://anses.hal.science/anses-05126539

Smith, A. F., Shinkins, B., Hall, P. S., Hulme, C. T., & Messenger, M. P. (2019). Toward a Framework for Outcome-Based Analytical Performance Specifications: A Methodology Review of Indirect Methods for Evaluating the Impact of Measurement Uncertainty on Clinical Outcomes. Clinical Chemistry, 65(11), 1363–1374. 10.1373/clinchem.2018.300954

Suchanek, M., & Robouch, P. (2009). Measurement uncertainty of test kit results – the ELISA example. Clinical Chemistry and Laboratory Medicine, 47(7), 808–810. 10.1515/CCLM.2009.181

van Leeuwen, C. C. E., Mulder, V. L., Batjes, N. H., & Heuvelink, G. B. M. (2022). Statistical modelling of measurement error in wet chemistry soil data. European Journal of Soil Science, 73(1), e13137. 10.1111/ejss.13137

Waugh, C. (2021). Factors affecting test reproducibility among laboratories: -EN--FR-Les facteurs affectant la reproductibilité des tests dans différents laboratoires -ES-Factores que inciden en la reproducibilidad de las pruebas entre laboratorios. Revue Scientifique et Technique de l’OIE, 40(1). 10.20506/rst.40.1.3213

WOAH. (2024). Measurement uncertainty. In Manual of Diagnostic Tests and Vaccines for Terrestrial Animals. https://www.woah.org/fileadmin/Home/eng/Health_standards/tahm/202405_Chapter_2.2.04_MEASUREMENT_UNCERT.pdf

