## Supplementary material for "Assessing measurement uncertainty at laboratory network scale and its impact on diagnostic performance in the absence of a gold standard: application to ELISA tests for *Coxiella burnetii in ruminants*": code and data: 1.outlier_detection.R

```
# loading packages
library(lme4)
library(tidyr)
library(dplyr)
library(ggplot2)


# loading ILTP data
source("./data/ILPT_data.R")


# running of the mixed model (takes a few minutes)

len <- Nlab * Nspecies * 2

test <- character(length = len)
serum <- character(length = len)
lab <- character(length = len)
type <- character(length = len)
sd <- numeric(length = len)
q2.5 <- numeric(length = len)
q97.5 <- numeric(length = len)

i <- 1

for (t in c("test1", "test2", "test3")) {
  
  labs <- sort(unique(subset(ILPT, test == t)$lab)) # laboratories that used test t
  
  for (l in labs) {
    
    for (s in c("cattle", "goat", "sheep")) {
      
      sub_data <- subset(tidy_ILPT, test == t & serum == s & lab != l)
      
      model <- lmer(ODR ~ batch + (1 | lab), data = sub_data)
      ci <- confint(model)
      
      test[i:(i+1)] <- rep(t, 2)
      serum[i:(i+1)] <- rep(s, 2)
      lab[i:(i+1)] <- rep(l, 2)
      type[i:(i+1)] <- c("inter", "intra")
      
      # inter- and intra-laboratory standard deviations and their confident interval
      sd[i:(i+1)] <- c(sqrt(VarCorr(model)$lab), sigma(model))
      q2.5[i:(i+1)] <- ci[1:2, 1]
      q97.5[i:(i+1)] <- ci[1:2, 2]
      
      i <- i+2
    }
  }
}

sd_l1o <- data.frame(test, serum, lab, type, sd, q2.5, q97.5) |>
  group_by(test, serum, type) |>
  mutate(med_q2.5 = median(q2.5), med_q97.5 = median(q97.5), med_sd = median(sd))


# parameters for graph aesthetic

Nlab_t1 <- length(unique(ILPT$lab[ILPT$test == "test1"]))
Nlab_t2 <- length(unique(ILPT$lab[ILPT$test == "test2"]))
Nlab_t3 <- length(unique(ILPT$lab[ILPT$test == "test3"]))
  
sd_l1o$xmin <- ifelse(sd_l1o$test == "test1", 0, 
                            ifelse(sd_l1o$test == "test2", Nlab_t1 + 1, Nlab_t1 + Nlab_t2 + 2))
sd_l1o$xmax <- ifelse(sd_l1o$test == "test1", Nlab_t1, 
                            ifelse(sd_l1o$test == "test2", 
                                   Nlab_t1 + Nlab_t2 + 1, Nlab_t1 + Nlab_t2 + Nlab_t3 + 2))
sd_l1o <- group_by(sd_l1o, test, serum, type) |>
  mutate(x = 1:n() + xmin)


# Figure A1.2

jpeg("./results/figA1.2.jpg", width = 1800, height = 1300)

ggplot(data = arrange(sd_l1o, lab)) +
  geom_rect(aes(ymin = med_q2.5, ymax = med_q97.5, xmin = xmin, xmax = xmax, fill = test), 
            alpha = 0.02) +
  geom_pointrange(aes(x = x, y = sd, ymin = q2.5, ymax = q97.5, color = test),
                  size = 0.8, linewidth = 0.8) +
  geom_text(aes(x = x, y = ifelse(sd < med_q2.5 | sd > med_q97.5, sd, NA), label = lab), 
            size = 10) +
  facet_grid(type ~ serum, scales = "free_y",
             labeller = labeller(type = c("inter" = "inter-laboratory",
                                          "intra" = "intra-laboratory"))) +
  scale_x_continuous(label = c("test1","test2","test3"), 
                     breaks = c(Nlab_t1/2, Nlab_t1 + Nlab_t2/2 + 1, Nlab_t1 + Nlab_t2 + Nlab_t3/2 + 2)) +
  labs(x = "", y = "standard deviation") +
  theme_bw() +
  theme(text = element_text(size = 40), legend.position = "none")

dev.off()


# outlier laboratories
out_lab <- unique(sd_l1o$lab[sd_l1o$sd < sd_l1o$med_q2.5 | sd_l1o$sd > sd_l1o$med_q97.5])

write.table(out_lab, "./intermediate_results/outlier_labs.csv", quote = FALSE)
```
