## Supplementary material for "Assessing measurement uncertainty at laboratory network scale and its impact on diagnostic performance in the absence of a gold standard: application to ELISA tests for *Coxiella burnetii in ruminants*": code and data: 2.measurement_uncertainty_assessment.R

```
# loading packages
library(tidyr)
library(rjags)
library(runjags)
library(lattice)
library(dplyr)
library(ggplot2)

# loading data
source("./data/ILPT_data_without_outlier.R")
source("./data/cut_offs.R")


set.seed(3)


# mixed model in JAGS language

mixed_model <- "model {

  for (i in 1:Nsample) {
    x[i] ~ dnorm(x0 + 
            L[numlab[i]] + 
            ifelse(numbatch[i] == Nbatch, -sum(b), b[min(numbatch[i], Nbatch - 1)]), 
            1 / Sintra^2)
            
    # the reference for the batch effect is the mean ODR measured across all batches,
    # so the sum of all the batch effects = 0, so b[Nbatch] = -sum(b)
  }
  
  for (j in 1:Nlab) {
    L[j] ~ dnorm(0, 1 / Sinter^2)
  }

  # priors

  for (k in 1:(Nbatch-1)) {
    b[k] ~ dnorm(0, 1 / 12.4^2)
  }

  x0 ~ dnorm(mu, 1 / 20^2)
  Sinter ~ dt(0, 1/20^2, 1)T(0,)
  Sintra ~ dt(0, 1/20^2, 1)T(0,)
} "


# data.frames to save the estimated parameters
df_x0_sd <- data.frame(matrix(ncol = 9, nrow = Ntest * Nspecies))
names(df_x0_sd) <- c("test", "serum", "x0", 
                     "Sinter", "Sinter_q1", "Sinter_q2", 
                     "Sintra", "Sintra_q1", "Sintra_q2")

df_batch_eff <- data.frame(matrix(ncol = 4, nrow = Nbatch * Nspecies))
names(df_batch_eff) <- c("serum", "test", "batch", "batch_eff")


# running of the mixed model for each serum and test (takes a few minutes)

i <- 1
j <- 0

for (s in c("cattle", "goat", "sheep")) {
  for (t in c("test1", "test2", "test3")) {
    
    # running of MCM model
    
    data <- subset(tidy_ILPT_out, serum == s & test == t)
    mu <- subset(tidy_mean_rnl, serum == s & test == t)$ODR
    
    Nlab_t <- length(unique(data$lab))
    
    test_batch <- unique(data$batch)
    Nbatch_t <- length(test_batch)
    if (Nbatch_t == 2) {batch_names <- "b"
    } else {batch_names <- paste0("b[",1:(Nbatch_t-1),"]")
    }
    
    data4jags <- list(Nlab = Nlab_t,
                      Nbatch = Nbatch_t,
                      Nsample = nrow(data),
                      numlab = match(data$lab, unique(data$lab)),
                      numbatch = match(data$batch, unique(data$batch)),
                      mu = mu,
                      x = data$ODR)
    
    inits <- list(list(Sintra = 2, Sinter = 5, x0 = 20, b = runif(Nbatch_t-1, -10, 10)),
                  list(Sintra = 5, Sinter = 10, x0 = 50, b = runif(Nbatch_t-1, -10, 10)),
                  list(Sintra = 10, Sinter = 20, x0 = 80, b = runif(Nbatch_t-1, -10, 10)))
    
    model <- jags.model(file = textConnection(mixed_model),
                        data = data4jags,
                        inits = inits,
                        n.chains = 3)
    
    update(model, 10000)
    
    mcmc <- coda.samples(model, c("Sintra", "Sinter", "x0", "b", "L"), n.iter = 100000, thin = 15)
    
    
    # checking of the convergence
    
    print(xyplot((mcmc), layout=c(3,6), main = paste(s, t, sep = " ")))
    print(gelman.diag(mcmc))
    
    
    # saving the posterior distributions
    
    write.table(as.matrix(mcmc)[, "Sinter"], 
                paste("./intermediate_results/post_distrib_Sinter", s, t, ".csv", 
                      sep = "_"), 
                quote = F)
    write.table(as.matrix(mcmc)[, "Sintra"], 
                paste("./intermediate_results/post_distrib_Sintra", s, t, ".csv", 
                      sep = "_"),
                quote = F)
    write.table(combine.mcmc(mcmc, vars = "L", collapse.chains = TRUE),
                paste("./intermediate_results/post_distrib_lab_eff", s, t, ".csv", 
                      sep = "_"),
                quote = F)
    write.table(combine.mcmc(mcmc, vars = "b", collapse.chains = TRUE), 
                paste("./intermediate_results/post_distrib_batch_eff", s, t, ".csv",
                      sep = "_"),
                quote = F)
        
    
    # filling the data frame containing the intercept and standard deviations
    
    quant <- summary(mcmc)$quantiles

    df_x0_sd$test[i] <- t
    df_x0_sd$serum[i] <- s
    df_x0_sd$x0[i] <- quant[["x0","50%"]]
    df_x0_sd[i, c("Sinter", "Sinter_q1", "Sinter_q2")] <- quant["Sinter", c("50%", "2.5%", "97.5%")]
    df_x0_sd[i, c("Sintra", "Sintra_q1", "Sintra_q2")] <- quant["Sintra", c("50%", "2.5%", "97.5%")]
    
    
    # filling the data frame containing the batch effects

    df_batch_eff$test[(j+1):(j+Nbatch_t)] <- t
    df_batch_eff$serum[(j+1):(j+Nbatch_t)] <- s
    df_batch_eff$batch[(j+1):(j+Nbatch_t)] <- as.character(test_batch)
    
    df_batch_eff$batch_eff[(j+1):(j+Nbatch_t-1)] <- quant[batch_names, "50%"]
    df_batch_eff$batch_eff[j+Nbatch_t] <- - sum(quant[batch_names, "50%"])
    
    
    i <- i + 1
    j <- j + Nbatch_t
  }
}

write.table(df_x0_sd, "./intermediate_results/mixed_model_x0_sd.csv")


# Figure 3

tidy_Sinter <- df_x0_sd[, c("test", "x0", "Sinter", "Sinter_q1", "Sinter_q2", "serum")] |>
  mutate(sd = "sigma inter")
names(tidy_Sinter) <- c("test", "x0", "med", "q1", "q2", "serum", "sd")

tidy_Sintra <- df_x0_sd[, c("test", "x0", "Sintra", "Sintra_q1", "Sintra_q2", "serum")] |>
  mutate(sd = "sigma intra")
names(tidy_Sintra) <- names(tidy_Sinter)

tidy_Sinter_Sintra <- rbind(tidy_Sinter, tidy_Sintra)

tidy_opt_co$serum <- tidy_opt_co$species


jpeg("./results/fig3.jpg", width = 1800, height = 1800)

ggplot() +
  geom_pointrange(data = tidy_Sinter_Sintra, size = 3.6, linewidth = 2.8,
                  aes(x = x0, y = med, ymin = q1, ymax = q2, color = serum)) +
  geom_vline(data = tidy_opt_co, aes(xintercept = co, color = serum), 
             linetype = "dashed", linewidth = 2.8) +
  scale_color_brewer(palette = "Dark2") +
  facet_grid(test ~ sd, labeller = labeller(sd = c("sigma inter" = "inter-laboratory",
                                                   "sigma intra" = "intra-laboratory"))) +
  labs(x = "ODR", y = "standard deviation", color = "") +
  theme_bw(base_size = 18) +
  theme(text = element_text(size = 65), strip.text = element_text(size = 65),
        axis.text = element_text(size = 65),
        legend.text = element_text(size = 65), legend.position = "bottom")

dev.off()


# Figure A5

df_batch_eff <- left_join(df_batch_eff, df_x0_sd[, c("test", "serum", "x0")]) |>
  mutate(mean_batch = x0 + batch_eff)
df_batch_eff$batch <- factor(df_batch_eff$batch, levels = unlist(batches_test))

batch_pal <- c("#E69F00", "#6A3D9A","#009E73","#56B4E9",
                        "#0072B2","#D55E00","#CC79A7", "#999933", "#117733", "#882255",
                        "#44AA99", "#AA4499")

jpeg("./results/figA5.jpg", width = 1700, height = 1700)

ggplot() +
  geom_point(data = tidy_ILPT_out, 
             aes(x = lab, y = ODR, color = batch), size = 4) +
  scale_color_manual(values = batch_pal) +
  labs(x = "laboratory", y = "ODR") +
  geom_hline(data = df_batch_eff, aes(yintercept = mean_batch, color = batch), linewidth = 1.4) +
  geom_hline(data = tidy_opt_co, aes(yintercept = co), linetype = "dashed", linewidth = 1.4) +
  geom_hline(data = df_x0_sd, aes(yintercept = x0), linewidth = 1.4) +
  facet_grid(serum ~ test, scale = "free") +
  theme_bw() +
  theme(axis.text.x = element_blank(), panel.grid.major.x = element_line(colour = "darkgrey"),
        text = element_text(size = 40), legend.position = "none")

dev.off()
```
