## Supplementary material for "Assessing measurement uncertainty at laboratory network scale and its impact on diagnostic performance in the absence of a gold standard: application to ELISA tests for *Coxiella burnetii in ruminants*": code and data: 3.changing_result_probability.R

```
# loading packages
library(tidyr)
library(dplyr)
library(cubature)
library(ggplot2)

# loading data
source("./data/kiteval4500_data.R")
source("./data/integrate_function.R")
df_x0_sd <- read.table("./intermediate_results/mixed_model_x0_sd.csv", header = TRUE)


species <- character(length = 9)
test <- character(length = 9)
xmin <- numeric(length = 9)
xmax <- numeric(length = 9)
prop <- numeric(length = 9)

i <- 1

for (s in c("cattle", "goat", "sheep")) {
  for (t in c("test1", "test2", "test3")) {
    
    sub_kiteval <- tidy_kiteval[tidy_kiteval$species == s & tidy_kiteval$test == t, ]
    
    # cut-off of test t in species s
    co <- tidy_opt_co[tidy_opt_co$species == s & tidy_opt_co$test == t, ]$co
    
    unique_ODR <- unique(sub_kiteval$ODR)
    ODR <- unique_ODR[unique_ODR < co] # ODRs of negative individuals
    
    # probability of being positive when retested
    p <- 1 - sapply(ODR, function(x) {
      hcubature(intg, -Inf, Inf,
                sigma_inter = df_x0_sd[df_x0_sd$serum == s & df_x0_sd$test == t, ]$Sinter,
                sigma_intra = df_x0_sd[df_x0_sd$serum == s & df_x0_sd$test == t, ]$Sintra,
                xobs = x, co = co)$integral})
    
    # ODR values for which the probability of changing result is > 5%
    xmin[i] <- ODR[which.min(abs(p - 0.05))]
    xmax[i] <- 2*co - xmin[i]
    
    # proportion of individuals who have a probability of changing result >5%
    prop[i] <- sum(sub_kiteval$ODR < xmax[i] & sub_kiteval$ODR > xmin[i]) / nrow(sub_kiteval)
    
    species[i] <- s
    test[i] <- t
    
    i <- i + 1
  }
}
df_change_res <- data.frame(species, test, xmin, xmax, prop)


# Figure 4

jpeg("./results/fig4.jpg", width = 1800, height = 1150)

ggplot(data = df_change_res) +
  geom_histogram(data = tidy_kiteval, aes(x = ODR), binwidth = 2) +
  geom_vline(aes(xintercept = xmin), color = "red", linewidth = 1) +
  geom_vline(aes(xintercept = xmax), color = "red", linewidth = 1) +
  geom_vline(data = tidy_opt_co, aes(xintercept = co), linetype = "dashed", linewidth = 1) +
  geom_text(data = df_change_res, aes(label = paste0(round(prop*100, 1),"%"), 
                                      x = 250, y = 130),
            size = 12) +
  facet_grid(species ~ test) +
  labs(y = "count")+
  theme_bw() +
  theme(text = element_text(size = 35), strip.text = element_text(size = 35))

dev.off()
```
