## Supplementary material for "Assessing measurement uncertainty at laboratory network scale and its impact on diagnostic performance in the absence of a gold standard: application to ELISA tests for *Coxiella burnetii in ruminants*": code and data: 4.PPP_calculation.R

```
# loading packages
library(tidyr)
library(dplyr)
library(rjags)
library(runjags)
library(lattice)

# loading data and model
source("./data/kiteval4500_data.R")


# creating data for the latent class model
# data frame with the number of individuals for each combination of the 3 test results in each herd

data_lcm <- group_by(.data = kiteval, species, herd, res, dpt, Nherd) |>
  summarise(nb = n()) |>
  pivot_wider(names_from = "res", values_from = "nb", values_fill = 0) |>
  mutate(dpt = as.numeric(as.factor(dpt)))

data_lcm <- data_lcm[, c("species", "dpt", "herd", "Nherd", as.character(1:8))]


# Model in JAGS language (adapted from Lurier et al.2021)

model.lclass <-  "model {
  # Number of animal from each population (herd) in each of the 8 categories of tests results
  for (i in 1:Npop) {
  
    n[i,] ~ dmulti(p[i,],N[i])
  
    p[i,1] <- Pinf[i] * ((1 - Se1) * (1 - Se2) * (1 - Se3) + covse_000) + 
      (1 - Pinf[i]) * ((Sp1 * Sp2 * Sp3) + covsp_000)
    p[i,5] <- Pinf[i] * ((1 - Se1) * (1 - Se2) * Se3  + covse_001) + 
      (1 - Pinf[i]) * (Sp1 * Sp2 * (1 - Sp3) + covsp_001)
    p[i,3] <- Pinf[i] * ((1 - Se1) * Se2 * (1 - Se3)  + covse_010 ) + 
      (1 - Pinf[i]) * (Sp1 * (1 - Sp2) * Sp3 + covsp_010)
    p[i,7] <- Pinf[i] * ((1 - Se1) * Se2 * Se3  + covse_011) + 
      (1 - Pinf[i]) * (Sp1 * (1 - Sp2) * (1 - Sp3) + covsp_011)
    p[i,2] <- Pinf[i] * (Se1 * (1 - Se2) * (1 - Se3)  + covse_100) + 
      (1 - Pinf[i]) * ((1 - Sp1) * Sp2 * Sp3 + covsp_100)
    p[i,6] <- Pinf[i] * (Se1 * (1 - Se2) * Se3  + covse_101) + 
      (1 - Pinf[i]) * ((1 - Sp1) * Sp2 * (1 - Sp3) + covsp_101)
    p[i,4] <- Pinf[i] * (Se1 * Se2 * (1 - Se3)  + covse_110) + 
      (1 - Pinf[i]) * ((1 - Sp1) * (1 - Sp2) * Sp3 + covsp_110)
    p[i,8] <- Pinf[i] * (Se1 * Se2 * Se3  + covse_111) + 
      (1 - Pinf[i]) * ((1 - Sp1) * (1 - Sp2) * (1 - Sp3) + covsp_111)
    }
  
  
  # zero inflated beta-binomial distribution of the prevalence in each heard
  
  for(e in 1:Npop) {
    herdstatus[e] ~ dbern(Pherd[numdpt[e]])
    P[e] ~ dbeta(mup * (1 - gamma) / gamma, (1 - mup) * (1 - gamma) / gamma) T(0.00001,0.99999)
    Pinf[e] <- P[e] * herdstatus[e]
  
    # P[e] is at least 1 out the herdsize
    y_P[e] ~ dinterval(P[e], 1/Nherd[e])
  }
  
  
  # Calculation of the covariance terms
  
  covse_100 <- covse_011 + covse_111 - covse_000
	covse_101 <- -(covse_001 + covse_011 + covse_111)
	covse_110 <- covse_000 + covse_001 - covse_111
	covse_010 <- -(covse_000 + covse_001 + covse_011)
	
	covsp_100 <- covsp_011 + covsp_111 - covsp_000
	covsp_101 <- -(covsp_001 + covsp_011 + covsp_111)
	covsp_110 <- covsp_000 + covsp_001 - covsp_111
	covsp_010 <- -(covsp_000 + covsp_001 + covsp_011)
	
	
	# Calculation of the PPP
	
	for (i in 1:Npop) {
	
		#for T1- T2- T3-
	  se_ppp[1,i] <- Pherd[numdpt[i]] * P[i] * ((1 - Se1) * (1 - Se2) * (1 - Se3) + covse_000)
	  sp_ppp[1,i] <- (1 - Pherd[numdpt[i]] * P[i]) * ((Sp1 * Sp2 * Sp3) + covsp_000)
	  
	  #for T1+ T2- T3-
	  se_ppp[2,i] <- Pherd[numdpt[i]] * P[i] * (Se1 * (1 - Se2) * (1 - Se3)  + covse_100)
	  sp_ppp[2,i] <- (1 - Pherd[numdpt[i]] * P[i]) * ((1 - Sp1) * Sp2 * Sp3 + covsp_100)
	  
	  #for T1- T2+ T3-
	  se_ppp[3,i] <- Pherd[numdpt[i]] * P[i] * ((1 - Se1) * Se2 * (1 - Se3)  + covse_010 )
	  sp_ppp[3,i] <- (1 - Pherd[numdpt[i]] * P[i]) * (Sp1 * (1 - Sp2) * Sp3 + covsp_010)
	  
	  #for T1+ T2+ T3-
	  se_ppp[4,i] <- Pherd[numdpt[i]] * P[i] * (Se1 * Se2 * (1 - Se3) + covse_110)
	  sp_ppp[4,i] <- (1 - Pherd[numdpt[i]] * P[i]) * ((1 - Sp1) * (1 - Sp2) * Sp3 + covsp_110)
	  
	  #for T1- T2- T3+
	  se_ppp[5,i] <- Pherd[numdpt[i]] * P[i] * ((1 - Se1) * (1 - Se2) * Se3  + covse_001)
	  sp_ppp[5,i] <- (1 - Pherd[numdpt[i]] * P[i]) * (Sp1 * Sp2 * (1 - Sp3) + covsp_001)
	  
	  #for T1+ T2- T3+
	  se_ppp[6,i] <- Pherd[numdpt[i]] * P[i] * (Se1 * (1 - Se2) * Se3  + covse_101)
	  sp_ppp[6,i] <- (1 - Pherd[numdpt[i]] * P[i]) * ((1 - Sp1) * Sp2 * (1 - Sp3) + covsp_101)
	  
	  #for T1- T2+ T3+
	  se_ppp[7,i] <- Pherd[numdpt[i]] * P[i] * ((1 - Se1) * Se2 * Se3  + covse_011)
	  sp_ppp[7,i] <- (1 - Pherd[numdpt[i]] * P[i]) * (Sp1 * (1 - Sp2) * (1 - Sp3) + covsp_011)
	  
	  #for T1+ T2+ T3+
	  se_ppp[8,i] <- Pherd[numdpt[i]] * P[i] * (Se1 * Se2 * Se3  + covse_111)
	  sp_ppp[8,i] <- (1 - Pherd[numdpt[i]] * P[i]) * ((1 - Sp1) * (1 - Sp2) * (1 - Sp3) + covsp_111)
	  
	  
	  for(t in 1:8) {
			# The probability of being positive given this combination of test results
			# (simple application of Bayes theorem anagolous to PPV = TP / (TP + FP))
			ppp[t,i] <- se_ppp[t,i] / (se_ppp[t,i] + sp_ppp[t,i])
	  }
	}
		
	
  #########
  # Prior #
  #########
  
  # Between herd prevalence of the jth departement 
  
  for(j in 1:Ndpt) {
    Pherd[j] ~ dbeta(0.5, 0.5)
  }

  mup ~ dbeta(0.5, 0.5) T(0.00001, 0.99999)
  gamma ~ dbeta(0.5, 0.5) T(0.00001, 0.99999)
  
  Se1 ~ dbeta(0.5, 0.5)
  Se2 ~ dbeta(0.5, 0.5)
  Se3 ~ dbeta(0.5, 0.5)
  Sp1 ~ dbeta(0.5, 0.5)
  Sp2 ~ dbeta(0.5, 0.5)
  Sp3 ~ dbeta(0.5, 0.5)

  covse_111 ~ dt(0, 1/0.039^2, 1)
  covse_011 ~ dt(0, 1/0.039^2, 1)
  covse_000 ~ dt(0, 1/0.039^2, 1)
  covse_001 ~ dt(0, 1/0.039^2, 1)
  
  covsp_111 ~ dt(0, 1/0.039^2, 1)
  covsp_011 ~ dt(0, 1/0.039^2, 1)
  covsp_000 ~ dt(0, 1/0.039^2, 1)
  covsp_001 ~ dt(0, 1/0.039^2, 1)
  

  ##########################
  # Inequality constraints #
  ##########################

  # Se > 1-Sp
  yse1 ~ dinterval(Se1, c(1-Sp1,1))
  yse2 ~ dinterval(Se2, c(1-Sp2,1))
  yse3 ~ dinterval(Se3, c(1-Sp3,1))
  
  # inequality constraints for covariance terms
  
  yse_000 ~ dinterval(covse_000, 
    c(-(1-Se1)*(1-Se2)*(1-Se3), min(1-Se1, min(1-Se2, 1-Se3)) - (1-Se1)*(1-Se2)*(1-Se3)))
  yse_001 ~ dinterval(covse_001, 
    c(-(1-Se1)*(1-Se2)*Se3, min(1-Se1, min(1-Se2, Se3)) - (1-Se1)*(1-Se2)*Se3))
  yse_010 ~ dinterval(covse_010, 
    c(-(1-Se1)*Se2*(1-Se3), min(1-Se1, min(Se2, 1-Se3)) - (1-Se1)*Se2*(1-Se3)))
  yse_011 ~ dinterval(covse_011, 
    c(-(1-Se1)*Se2*Se3, min(1-Se1, min(Se2, Se3)) - (1-Se1)*Se2*Se3))
  yse_100 ~ dinterval(covse_100, 
    c(-Se1*(1-Se2)*(1-Se3), min(Se1, min((1-Se2), (1-Se3))) - Se1*(1-Se2)*(1-Se3)))
  yse_101 ~ dinterval(covse_101, 
    c(-Se1*(1-Se2)*Se3, min(Se1, min((1-Se2), Se3)) - Se1*(1-Se2)*Se3))
  yse_110 ~ dinterval(covse_110, 
    c(-Se1*Se2*(1-Se3), min(Se1, min(Se2, 1-Se3)) - Se1*Se2*(1-Se3)))
  yse_111 ~ dinterval(covse_111,
    c(-Se1*Se2*Se3, min(Se1, min(Se2, Se3)) - Se1*Se2*Se3))
  
  ysp_000 ~ dinterval(covsp_000,
    c(-Sp1*Sp2*Sp3, min(Sp1, min(Sp2, Sp3)) - Sp1*Sp2*Sp3))
  ysp_001 ~ dinterval(covsp_001, 
    c(-Sp1*Sp2*(1-Sp3), min(Sp1, min(Sp2, 1-Sp3)) - Sp1*Sp2*(1-Sp3)))
  ysp_010 ~ dinterval(covsp_010, 
    c(-Sp1*(1-Sp2)*Sp3, min(Sp1, min(1-Sp2, Sp3)) - Sp1*(1-Sp2)*Sp3))
  ysp_011 ~ dinterval(covsp_011,
    c(-Sp1*(1-Sp2)*(1-Sp3), min(Sp1, min(1-Sp2, 1-Sp3)) - Sp1*(1-Sp2)*(1-Sp3)))
  ysp_100 ~ dinterval(covsp_100, 
    c(-(1-Sp1)*Sp2*Sp3, min((1-Sp1), min(Sp2, Sp3)) - (1-Sp1)*Sp2*Sp3))
  ysp_110 ~ dinterval(covsp_110, 
    c(-(1-Sp1)*(1-Sp2)*Sp3, min((1-Sp1), min((1-Sp2), Sp3)) - (1-Sp1)*(1-Sp2)*(1-Sp3)))
  ysp_101 ~ dinterval(covsp_101, 
    c(-(1-Sp1)*Sp2*(1-Sp3), min((1-Sp1), min(Sp2, 1-Sp3)) - (1-Sp1)*Sp2*(1-Sp3)))
  ysp_111 ~ dinterval(covsp_111,
    c(-(1-Sp1)*(1-Sp2)*(1-Sp3), min(1-Sp1, min(1-Sp2, 1-Sp3)) - (1-Sp1)*(1-Sp2)*(1-Sp3)))
} 
"
  
  
# Function that runs the model and saves the PPP posterior distributions
  
  # Inputs:
  # data_lcm: data frame with one line per herd,
  #   and in column: the species, the department id, the herd id, the number of
  #   individuals in the herd, and the number of individuals having each combination
  #   of the three test results
  # sp: "cattle", "goat" or "sheep"
  
  fun_lcm <- function(data_lcm, sp) {
    
    data <- arrange(.data = subset(data_lcm, species == sp), herd)
    data$Nherd[is.na(data$Nherd)] <- median(data$Nherd, na.rm = T)
    
    data4jags <- list(Npop = nrow(data),
                      N = rowSums(data[,5:12]),
                      Nherd = data$Nherd,
                      n = as.matrix(data[,5:12]),
                      Ndpt = 10,
                      numdpt = data$dpt,
                      yse_000 = 1,
                      yse_001 = 1,
                      yse_010 = 1,
                      yse_011 = 1,
                      yse_100 = 1,
                      yse_101 = 1,
                      yse_110 = 1,
                      yse_111 = 1,
                      ysp_000 = 1,
                      ysp_001 = 1,
                      ysp_010 = 1,
                      ysp_011 = 1,
                      ysp_100 = 1,
                      ysp_101 = 1,
                      ysp_110 = 1,
                      ysp_111 = 1,
                      yse1 = 1,
                      yse2 = 1,
                      yse3 = 1,
                      y_P = rep(1,nrow(data)))
    
    ##################
    # Initialisation #
    ##################
    
    # initialization ensures that the parameters respect the constraints of the model
    
    inits <- list(list(),list(),list())
    
    for (k in 1:3) {
      
      Sp1 <- rbeta(1,5,1)
      Sp2 <- rbeta(1,5,1)
      Sp3 <- rbeta(1,5,1)
      
      Se1 <- runif(1,1-Sp1,1)
      Se2 <- runif(1,1-Sp2,1)
      Se3 <- runif(1,1-Sp3,1)
      
      
      covse_111 <- 0
      covse_011 <- 0
      covse_001 <- 0
      covse_000 <- 0
      
      covsp_111 <- 0
      covsp_011 <- 0
      covsp_001 <- 0
      covsp_000 <- 0
      
      
      P <- runif(length(data$Nherd), 1/data$Nherd, 1)
      
      inits[[k]] <- list(Se1 = Se1, Se2 = Se2, Se3 = Se3,Sp1 = Sp1,
                         Sp2 = Sp2, Sp3 = Sp3, covse_111 = covse_111,
                         covse_011 = covse_011, covse_001 = covse_001,
                         covse_000 = covse_000, covsp_111 = covsp_111,
                         covsp_011 = covsp_011, covsp_001 = covsp_001,
                         covsp_000 = covsp_000, P = P)
    }
    
    m.lclass <- jags.model(file = textConnection(model.lclass), data = data4jags,  
                           n.chains = 3, inits = inits)
    
    update(m.lclass, n.iter = 10000) # burnin
    
    mcmc.lclass<- coda.samples(m.lclass, c("ppp"), 
                               n.iter = 100000, thin = 20) 
    
        write.table(combine.mcmc(mcmc.lclass, vars = "ppp", collapse.chains = T), 
                paste0("./intermediate_results/ppp_", sp, ".csv"), quote = F)
    
    
    # Check of the convergence 
    print(xyplot((mcmc.lclass), layout = c(3,6)))
  }


# running of the model

set.seed(3)

fun_lcm(data_lcm, "cattle") # takes approximatively 1 hour
fun_lcm(data_lcm, "goat") # takes approximatively 1 hour
fun_lcm(data_lcm, "sheep") # takes approximatively 1 hour
```
