## Supplementary material for "Assessing measurement uncertainty at laboratory network scale and its impact on diagnostic performance in the absence of a gold standard: application to ELISA tests for *Coxiella burnetii in ruminants*": code and data: 5.Se_Sp_estimation.R

```
# loading packages
library(tidyr)
library(dplyr)
library(parallel)
library(cubature)
library(ggplot2)
library(gridExtra)
library(ggtext)

#loading data
source("./data/kiteval4500_data.R")
source("./data/ILPT_data_without_outlier.R")
source("./data/integrate_function.R")


set.seed(3)


# This step takes several hours
# You can directly run the 'results' part below, which provides figures and table.
# To reduce the run time, you can decrease the number of iterations n_iter
# or run it for only one species and/or one test.


n_iter <- 10000

l1 <- Nspecies * Ntest * 2
l2 <- Nspecies * Nlab * 2
l3 <- Nspecies * Nbatch * 2

list_SeSp <- vector("list", l1)
list_SeSp_lab <- vector("list", l2)
list_SeSp_batch <- vector("list", l3)

i1 <- 1
i2 <- 1
i3 <- 1


cl <- makeCluster(detectCores()-1)


for (s in c("cattle", "goat", "sheep")) {
  
  #posterior distribution of the PPPs
  ppp_post <- read.table(paste0("./intermediate_results/ppp_", s, ".csv"), header = T)
  
  sub_kiteval1 <- tidy_kiteval[tidy_kiteval$species == s, ] |>
    mutate(herd2 = as.numeric(as.factor(herd)))
  
  for (t in c("test1", "test2", "test3")) {
    
    sub_kiteval <- sub_kiteval1[sub_kiteval1$test == t, ]
    
    sub_co <- tidy_opt_co[tidy_opt_co$species == s & tidy_opt_co$test == t,]$co # cut-off
    
    N <- nrow(sub_kiteval) # number of individuals
    
    # posterior distributions
    Sinter_post <- read.table(paste("./intermediate_results/post_distrib_Sinter",
                                    s, t, ".csv", sep = "_"), 
                              header = T)
    Sintra_post <- read.table(paste("./intermediate_results/post_distrib_Sintra", 
                                    s, t, ".csv", sep = "_")
                              , header = T)
    lab_eff_post <- read.table(paste("./intermediate_results/post_distrib_lab_eff", 
                                     s, t, ".csv", sep = "_"), 
                               header = T)
    batch_eff_post <- read.table(paste("./intermediate_results/post_distrib_batch_eff", 
                                       s, t, ".csv", sep = "_"), 
                               header = T)
    
    # random draw of the parameters in their posterior distributions
    sampled_indices <- sample(1:nrow(Sinter_post), n_iter, replace = TRUE)
    s_inter <- Sinter_post[sampled_indices,]
    s_intra <- Sintra_post[sampled_indices,]
    lab_eff <- lab_eff_post[sampled_indices,]
    batch_eff <- as.data.frame(batch_eff_post[sampled_indices,])
    
    
    ## PPP ##
    
    # PPPs from their posterior distribution 
    # (matrix with one line per iteration and one column per kiteval individual)
    ppp <- as.matrix(ppp_post[sample(1:nrow(ppp_post), n_iter),
                              paste("ppp", sub_kiteval$res, sub_kiteval$herd2, "" , sep = ".")])
    
    
    ### probability to be negative when retested ###
    
    unique_ODR <- unique(sub_kiteval$ODR)
    len_odr <- length(unique_ODR)

    clusterExport(cl, c("hcubature", "intg", "s_inter", "s_intra", 
                        "unique_ODR", "sub_co", "n_iter"))
    
    prob_neg_unique <- parSapply(cl, unique_ODR, function(x) {
      sapply(1:n_iter, function(j) {
        hcubature(intg, -Inf, Inf,
                  sigma_inter = s_inter[j], sigma_intra = s_intra[j],
                  xobs = x, co = sub_co)$integral
        })
      })
    # probability to be negative
    # (matrix with one line per iteration and one column per individual)
    prob_neg <- prob_neg_unique[, match(sub_kiteval$ODR, unique_ODR)]
    
    
    # vector of the PPP sum for all individuals
    sum_ppp <- rowSums(ppp)
    
    # vector of the sum of the probabilities to be negative for all individuals
    sum_prob_neg <- rowSums(prob_neg)
    
    # vector of the sum of PPP*prob_neg for all individuals
    sum_ppp_prob_neg <- rowSums(prob_neg * ppp)
    
    # vector of the number of observed negative individuals
    sum_neg <- sum(sub_kiteval$neg)
    
    # vector of the sum of PPP*neg for all individuals
    sum_ppp_neg <- rowSums(sweep(ppp, 2, sub_kiteval$neg, "*"))
    

    ## diagnostic performance accounting and not accounting for measurement uncertainty ##
    
    vec_se <- (sum_ppp - sum_ppp_neg) / sum_ppp
    vec_sp <- (sum_neg - sum_ppp_neg) / (N - sum_ppp)
    vec_se_u <- (sum_ppp - sum_ppp_prob_neg) / sum_ppp
    vec_sp_u <- (sum_prob_neg - sum_ppp_prob_neg) / (N - sum_ppp)

    vec_fp <- N - sum_ppp - sum_neg + sum_ppp_neg
    vec_fp_u <- N - sum_ppp - sum_prob_neg + sum_ppp_prob_neg
    
    
    list_SeSp[[i1]] <- list(species = s, 
                            test = t,
                            
                            med_Se = median(vec_se), 
                            med_Sp = median(vec_sp),
                            q1_Se = quantile(vec_se, probs = 0.025),
                            q2_Se = quantile(vec_se, probs = 0.975),
                            q1_Sp = quantile(vec_sp, probs = 0.025),
                            q2_Sp = quantile(vec_sp, probs = 0.975),
                            
                            med_Se_u = median(vec_se_u), 
                            med_Sp_u = median(vec_sp_u),
                            q1_Se_u = quantile(vec_se_u, probs = 0.025),
                            q2_Se_u = quantile(vec_se_u, probs = 0.975),
                            q1_Sp_u = quantile(vec_sp_u, probs = 0.025),
                            q2_Sp_u = quantile(vec_sp_u, probs = 0.975),

                            FP = median(vec_fp),
                            q1_FP = quantile(vec_fp, probs = 0.025),
                            q2_FP = quantile(vec_fp, probs = 0.975),
                            FN = median(sum_ppp_neg),
                            q1_FN = quantile(sum_ppp_neg, probs = 0.025),
                            q2_FN = quantile(sum_ppp_neg, probs = 0.975),
                            
                            FP_u = median(vec_fp_u),
                            q1_FP_u = quantile(vec_fp_u, probs = 0.025),
                            q2_FP_u = quantile(vec_fp_u, probs = 0.975),
                            FN_u = median(sum_ppp_prob_neg),
                            q1_FN_u = quantile(sum_ppp_prob_neg, probs = 0.025),
                            q2_FN_u = quantile(sum_ppp_prob_neg, probs = 0.975))
    i1 <- i1 + 1
    
    
    ## Se and Sp of each lab ##
    
    Nlab_t <- ncol(lab_eff)
    
    for (l in 1:Nlab_t) {
      
      # results (0 for positive, 1 for negative) in lab l 
      # (one line per iteration and one column per individual)
      res_lab <- outer(lab_eff[,l], sub_kiteval$ODR, "+") < sub_co

      # sum of the PPPs of the individuals negative in lab l
      sum_ppp_res_lab <- rowSums(res_lab * ppp)
      
      vec_se_lab <- (sum_ppp - sum_ppp_res_lab) / sum_ppp
      vec_sp_lab <- (rowSums(res_lab) - sum_ppp_res_lab) / (N - sum_ppp)
      
      
      list_SeSp_lab[[i2]] <- list(species = s,
                                  lab = unlist(labs_test[t])[l],
                                  Se = median(vec_se_lab),
                                  Sp = median(vec_sp_lab),
                                  q1_Se = quantile(vec_se_lab, 0.025),
                                  q1_Sp = quantile(vec_sp_lab, 0.025),
                                  q2_Se = quantile(vec_se_lab, 0.975),
                                  q2_Sp = quantile(vec_sp_lab, 0.975))
      i2 <- i2 + 1
    }
    
    
    ## Se and Sp of each batch ##
    
    Nbatch_t <- ncol(batch_eff) + 1
    batch_eff$b. <- apply(batch_eff, 1, function(x) {- sum(x)})
    
    for (b in 1:Nbatch_t) {
      
      # results (0 for positive, 1 for negative) with batch b 
      #(one line per iteration and one column per individual)
      res_batch <- outer(batch_eff[,b], sub_kiteval$ODR, "+") < sub_co

      # sum of the PPPs of the individuals negative with batch b
      sum_ppp_res_batch <- rowSums(res_batch * ppp)
      
      vec_se_batch <- (sum_ppp - sum_ppp_res_batch) / sum_ppp
      vec_sp_batch <- (rowSums(res_batch) - sum_ppp_res_batch) / (N - sum_ppp)
      
      
      list_SeSp_batch[[i3]] <- list(species = s,
                                    batch = unlist(batches_test[t])[b],
                                    Se = median(vec_se_batch),
                                    Sp = median(vec_sp_batch),
                                    q1_Se = quantile(vec_se_batch, 0.025),
                                    q1_Sp = quantile(vec_sp_batch, 0.025),
                                    q2_Se = quantile(vec_se_batch, 0.975),
                                    q2_Sp = quantile(vec_sp_batch, 0.975))
      
     i3 <- i3 + 1
    }
    
  }
}

stopCluster(cl)


# Se, Sp, FN and FP with and without measurement uncertainty
Se_Sp <- do.call(rbind, lapply(list_SeSp, as.data.frame))

# Se and Sp of each lab
Se_Sp_lab <- do.call(rbind, lapply(list_SeSp_lab, as.data.frame))

# Se and Sp of each batch
Se_Sp_batch <- do.call(rbind, lapply(list_SeSp_batch, as.data.frame))

write.table(Se_Sp, "./results/Se_Sp_u.csv", quote = F)
write.table(Se_Sp_lab, "./results/Se_Sp_lab.csv", quote = F)
write.table(Se_Sp_batch, "./results/Se_Sp_batch.csv", quote = F)


### Results ###

Se_Sp <- read.table("./intermediate_results/Se_Sp_u.csv", header = T)
Se_Sp_lab <- read.table("./intermediate_results/Se_Sp_lab.csv", header = T)
Se_Sp_batch <- read.table("./intermediate_results/Se_Sp_batch.csv", header = T)


# Figure 5

tidy_Se_Sp <- Se_Sp[, 1:14] |>
  pivot_longer(cols = -c(species, test)) |>
  mutate(u = grepl("_u$", name), 
         name = gsub("_u$", "", name)) |>
  separate(name, into = c("stat", "par"), sep = "_", fill = "right") |>
  pivot_wider(names_from = "stat", values_from = "value") |>
  mutate(u = factor(u, levels = c("FALSE", "TRUE"), labels = c("no", "yes")))


jpeg("./results/fig5.jpg", width = 1800, height = 1500)

ggplot(data = tidy_Se_Sp) +
  geom_pointrange(aes(x = species, y = med, ymin = q1, ymax = q2, color = u),
                  position = position_dodge(0.5), size = 3.6, linewidth = 2.8) +
  scale_color_discrete(name = "accounting for measurement uncertainty: ") +
  facet_grid(par ~ test, scale = "free", 
             labeller = labeller(par = c("Se"="sensitivity", "Sp"="specificity"))) +
  labs(x = "", y = "") +
  theme_bw(base_size = 18) +
  theme(text = element_text(size = 65), strip.text = element_text(size = 65), 
        axis.text = element_text(size = 58),
        legend.text = element_text(size = 65), legend.position = "bottom")

dev.off()


# Table A4

#number of FN and FP in kiteval data
FN_FP <- Se_Sp[, c(1,2, 15:26)]  |>
  left_join(N_per_species)

#number of FN and FP for 1000 individuals
FN_FP[3:14] <- round(FN_FP[3:14] * 1000 / FN_FP$N, 1)

FN_FP <- mutate(FN_FP,
                FN_without_u = paste0(FN, " [", q1_FN, "; ", q2_FN, "]"),
                FN_with_u = paste0(FN_u, " [", q1_FN_u, "; ", q2_FN_u, "]"),
                FP_without_u = paste0(FP, " [", q1_FP, "; ", q2_FP, "]"),
                FP_with_u = paste0(FP_u, " [", q1_FP_u, "; ", q2_FP_u, "]"))

tableA4 <- FN_FP[, c(1,2, 16:19)]

write.table(tableA4, "./results/tableA4.csv")


# Figure 6

# parameters for graph aesthetic

lab_min_max <- data.frame(test = c("test1", "test2", "test3"),
                          xmin = c(unlist(labs_test["test1"])[1], 
                                   unlist(labs_test["test2"])[1], 
                                   unlist(labs_test["test3"])[1]),
                          xmax = c(tail(unlist(labs_test["test1"]), 1),
                                   tail(unlist(labs_test["test2"]), 1),
                                   tail(unlist(labs_test["test3"]), 1)))

Se_Sp <- Se_Sp |>
  left_join(lab_min_max) |>
  mutate(xmin = factor(xmin, levels = unlist(labs_test)),
         xmax = factor(xmax, levels = unlist(labs_test)))

Se_Sp_lab$lab <- factor(Se_Sp_lab$lab, levels = unlist(labs_test))
Se_Sp_lab <- left_join(Se_Sp_lab, unique(ILPT[, c("lab", "test")]))


graph_Se_lab <- ggplot() +
  geom_pointrange(data = Se_Sp_lab, linewidth = 1.3, size = 0.9,
                  aes(x = lab, y = Se, ymin = q1_Se, ymax = q2_Se, color = test)) +
  geom_segment(data = Se_Sp, linewidth = 1.3,
               aes(x = xmin, xend = xmax, y = med_Se, yend = med_Se, color = test)) +
  geom_rect(data = Se_Sp, alpha = 0.3,
            aes(xmin = xmin, xmax = xmax, ymin = q1_Se, ymax = q2_Se, fill = test)) +
  facet_grid(species ~ .) +
  labs(x = "laboratory", y = "", title = "sensitivity") +
  theme_bw() +
  theme(text = element_text(size = 35), strip.text = element_text(size = 35),
        axis.text.x = element_blank(), axis.title.x = element_text(size = 35),
        legend.text = element_text(size = 35), legend.position = "none",
        panel.grid.major.x = element_line(colour = "darkgrey"),
        plot.title = element_textbox(fill = "grey85", box.colour = "grey20",
                                     linewidth = 0.5, linetype = "solid", 
                                     width = grid::unit(1, "npc"), halign = 0.5,
                                     size = 35)) +
  scale_y_continuous(breaks = c(0.60, 0.80, 1.00), labels = c("0.60", "0.80", "1.00"))

graph_Sp_lab <- ggplot() +
  geom_pointrange(data = Se_Sp_lab, linewidth = 1.3, size = 0.9,
                  aes(x = lab, y = Sp, ymin = q1_Sp, ymax = q2_Sp, color = test)) +
  geom_segment(data = Se_Sp, linewidth = 1.3,
               aes(x = xmin, xend = xmax, y = med_Sp, yend = med_Sp, color = test)) +
  geom_rect(data = Se_Sp, alpha = 0.3,
            aes(xmin = xmin, xmax = xmax, ymin = q1_Sp, ymax = q2_Sp, fill = test)) +
  scale_color_discrete(name = "") +
  scale_fill_discrete(name = "") +
  facet_grid(species ~ .) +
  labs(x = "laboratory", y = "", title = "specificity") +
  theme_bw() +
  theme(text = element_text(size = 35), strip.text = element_text(size = 35),
        axis.text.x = element_blank(), axis.title.x = element_text(size = 35),
        legend.text = element_text(size = 35), legend.position = "bottom",
        panel.grid.major.x = element_line(colour = "darkgrey"),
        plot.title = element_textbox(fill = "grey85", box.colour = "grey20",
                                     linewidth = 0.5, linetype = "solid", 
                                     width = grid::unit(1, "npc"), halign = 0.5,
                                     size = 35))


#getting legend
tmp <- ggplot_gtable(ggplot_build(graph_Sp_lab))
leg <- which(sapply(tmp$grobs, function(x) x$name) == "guide-box")
legend <- tmp$grobs[[leg]]


jpeg("./results/fig6.jpg", width = 1800, height = 1375)

grid.arrange(graph_Se_lab, graph_Sp_lab + theme(legend.position = "none"), legend, 
             ncol = 2, nrow = 2, layout_matrix= rbind(c(1,2), 3), heights = c(2.5, 0.15))

dev.off()


# Figure 7

batch_min_max <- data.frame(test = c("test1", "test2", "test3"),
                            xminb = c(unlist(batches_test["test1"])[1], 
                                      unlist(batches_test["test2"])[1], 
                                      unlist(batches_test["test3"])[1]),
                            xmaxb = c(tail(unlist(batches_test["test1"]), 1),
                                      tail(unlist(batches_test["test2"]), 1),
                                      tail(unlist(batches_test["test3"]), 1)))

Se_Sp <- Se_Sp |>
  left_join(batch_min_max) |>
  mutate(xmin = factor(xmin, levels = unlist(batches_test)),
         xmax = factor(xmax, levels = unlist(batches_test)))

Se_Sp_batch <- left_join(Se_Sp_batch, unique(ILPT[, c("batch", "test")]))
Se_Sp_batch$batch <- factor(rep(unlist(batches_test), 3), levels = unlist(batches_test))


graph_Se_batch <- ggplot() +
  geom_pointrange(data = Se_Sp_batch, linewidth = 2.8, size = 2.5,
                  aes(x = batch, y = Se, ymin = q1_Se, ymax = q2_Se, color = test)) +
  geom_segment(data = Se_Sp, linewidth = 2.8,
               aes(x = xminb, xend = xmaxb, y = med_Se, yend = med_Se, color = test)) +
  geom_rect(data = Se_Sp, alpha = 0.3,
            aes(xmin = xminb, xmax = xmaxb, ymin = q1_Se, ymax = q2_Se, fill = test)) +
  facet_grid(species ~ .) +
  labs(x = "batch", y = "", title = "sensitivity") +
  theme_bw(base_size = 18) +
  theme(axis.text.x = element_blank(), axis.title.x = element_text(size = 65),
        text = element_text(size = 65), strip.text = element_text(size = 65),
        panel.grid.major.x = element_line(colour = "darkgrey"),
        legend.text = element_text(size = 65), legend.position = "none",
        plot.title = element_textbox(fill = "grey85", box.colour = "grey20",
                                     linewidth = 0.5, linetype = "solid", 
                                     width = grid::unit(1, "npc"), halign = 0.5,
                                     size = 65))

graph_Sp_batch <- ggplot() +
  geom_pointrange(data = Se_Sp_batch, linewidth = 2.8, size = 2.5,
                  aes(x = batch, y = Sp, ymin = q1_Sp, ymax = q2_Sp, color = test)) +
  geom_segment(data = Se_Sp, linewidth = 2.8,
               aes(x = xminb, xend = xmaxb, y = med_Sp, yend = med_Sp, color = test)) +
  geom_rect(data = Se_Sp, alpha = 0.3,
            aes(xmin = xminb, xmax = xmaxb, ymin = q1_Sp, ymax = q2_Sp, fill = test)) +
  scale_color_discrete(name = "") +
  scale_fill_discrete(name = "") +
  facet_grid(species ~ .) +
  labs(x = "batch", y = "", title = "specificity") +
  theme_bw(base_size = 18) +
  theme(axis.text.x = element_blank(), axis.title.x = element_text(size = 65),
        text = element_text(size = 66), strip.text = element_text(size = 65),
        legend.text = element_text(size = 65), legend.position = "bottom",
        panel.grid.major.x = element_line(colour = "darkgrey"),
        plot.title = element_textbox(fill = "grey85", box.colour = "grey20",
                                     linewidth = 0.5, linetype = "solid", 
                                     width = grid::unit(1, "npc"), halign = 0.5,
                                     size = 65)) +
  scale_y_continuous(breaks = seq(0.92, 1, 0.04))


#getting legend
tmp_b <- ggplot_gtable(ggplot_build(graph_Sp_batch))
leg_b <- which(sapply(tmp_b$grobs, function(x) x$name) == "guide-box")
legend_b <- tmp_b$grobs[[leg_b]]


jpeg("./results/fig7.jpg", width = 1800, height = 1800)

grid.arrange(graph_Se_batch, graph_Sp_batch + theme(legend.position = "none"), legend_b,
             ncol = 2, nrow = 2, layout_matrix= rbind(c(1,2), 3), heights = c(2.5, 0.15))
dev.off()
```
