## Supplementary material for "Assessing measurement uncertainty at laboratory network scale and its impact on diagnostic performance in the absence of a gold standard: application to ELISA tests for *Coxiella burnetii in ruminants*": code and data: cut_offs.R

```
# cut-offs

opt_co <- data.frame(test1 = c(44.9, 6.6, 26.2),
                     test2 = c(33.2, 18.9, 55),
                     test3 = c(84.3, 49.9, 89.7),
                     species = c("cattle", "goat", "sheep"))

tidy_opt_co <- pivot_longer(opt_co, cols = c("test1", "test2", "test3"), 
                            names_to = "test", values_to = "co")
```
