## Supplementary material for "Assessing measurement uncertainty at laboratory network scale and its impact on diagnostic performance in the absence of a gold standard: application to ELISA tests for *Coxiella burnetii in ruminants*": code and data: ILPT_data.R

```
ILPT <- read.table("./data/ILPT.csv", header = T, dec = ",", sep = ";")

colnames(ILPT)[1:3] <- c("lab", "test", "batch")

ILPT$test [ILPT$test == "ID Screen\xb8 Q fever indirect"] <- "test3"
ILPT$test [ILPT$test == "BioX MonoscreenAbELISA Coxiella blocking phase I (BIO K 001)"] <- "test4"
ILPT$test [ILPT$test == "IDEXX Q Feber Ab Test"] <- "test1"
ILPT$test [ILPT$test == "PrioCHECK? Ruminant Q Fever Ab Plate Kit"] <- "test2"

ILPT$lab <- as.factor(ILPT$lab)
ILPT$test <- as.factor(ILPT$test)
ILPT$batch <- as.factor(ILPT$batch)


# selection of the tests and sera used in our study
ILPT <- ILPT[ILPT$test != "test4", 
             !grepl("B1", names(ILPT)) & !(names(ILPT) %in% c("N1", "N2", "P1", "P2"))]


tidy_ILPT <- pivot_longer(data = ILPT, 
                          cols = c("B2R1", "B2R2", "B2R3", "B2R4", 
                                   "CR1", "CR2", "CR3", "CR4", 
                                   "OR1", "OR2", "OR3", "OR4"), 
                          names_to = "Replicate",
                          values_to = "ODR")

tidy_ILPT$serum <- factor(gsub("R[1-4]", "", tidy_ILPT$Replicate), 
                          levels = c("B2", "C", "O"),
                          labels = c("cattle", "goat", "sheep"))


Ntest <- length(unique(ILPT$test)) # number of tests
Nspecies <- length(unique(tidy_ILPT$serum)) # number of species
Nlab <- length(unique(ILPT$lab)) # number of labs
```
