## Supplementary material for "Assessing measurement uncertainty at laboratory network scale and its impact on diagnostic performance in the absence of a gold standard: application to ELISA tests for *Coxiella burnetii in ruminants*": code and data: ILPT_data_without_outlier.R

```
source("./data/ILPT_data.R")

# outlier laboratories
out_lab <- as.vector(read.table("./intermediate_results/outlier_labs.csv", header = TRUE))$x

# data without outlier
tidy_ILPT_out <- tidy_ILPT[tidy_ILPT$lab %in% out_lab == FALSE, ]


Nlab <- length(unique(tidy_ILPT_out$lab)) # number of labs
Nbatch <- length(unique(tidy_ILPT_out$batch)) # number of batches


# batches used by each test
batches_test <- list(test1 = unique(tidy_ILPT_out[tidy_ILPT_out$test == "test1", "batch"]),
                     test2 = unique(tidy_ILPT_out[tidy_ILPT_out$test == "test2", "batch"]),
                     test3 = unique(tidy_ILPT_out[tidy_ILPT_out$test == "test3", "batch"]))

# sorting the batches in test order
tidy_ILPT_out$batch <- factor(tidy_ILPT_out$batch, levels = unlist(batches_test))


# laboratories used by each test
labs_test <- list(test1 = unique(tidy_ILPT_out[tidy_ILPT_out$test == "test1", "lab"]),
                  test2 = unique(tidy_ILPT_out[tidy_ILPT_out$test == "test2", "lab"]),
                  test3 = unique(tidy_ILPT_out[tidy_ILPT_out$test == "test3", "lab"]))


# mean ODR of the sera measured by the French Reference Laboratory
mean_rnl <- data.frame(serum = c("cattle", "goat", "sheep"),
                       test1 = c(39.37, 9.58, 43.24),
                       test2 = c(22.33, 11.23, 26.65),
                       test3 = c(84.54, 39.89, 73.51))

tidy_mean_rnl <- pivot_longer(mean_rnl, cols = c("test1", "test2", "test3"), 
                              names_to = "test", values_to = "ODR")
```
