## Supplementary material for "Assessing measurement uncertainty at laboratory network scale and its impact on diagnostic performance in the absence of a gold standard: application to ELISA tests for *Coxiella burnetii in ruminants*": code and data: integrate_function.R

```
# function to be integrated for the calculation of the probability of changing result when retested

intg <- function(u, xobs, sigma_inter, sigma_intra, co) {
  return(dnorm(u, xobs, sigma_inter) * pnorm(co, u, sigma_intra))
}
```
