## Supplementary material for "Assessing measurement uncertainty at laboratory network scale and its impact on diagnostic performance in the absence of a gold standard: application to ELISA tests for *Coxiella burnetii in ruminants*": code and data: kiteval4500_data.R

```
source("./data/cut_offs.R")

kiteval <- read.table("./data/kiteval4500.csv", header = TRUE)

kiteval <- left_join(kiteval, opt_co, by = "species", suffix = c("","_co")) |>
  mutate(res1 = as.numeric(test1 >= test1_co), # result with test 1
         res2 = as.numeric(test2 >= test2_co), # result with test 2
         res3 = as.numeric(test3 >= test3_co), # result with test 3
         res = as.factor(as.numeric(interaction(res1, res2, res3)))) # combination of the 3 test results

tidy_kiteval <- pivot_longer(data = kiteval[,c("dpt", "herd", "species", 
                                               "test1", "test2", "test3", "res")], 
                             cols = c("test1", "test2", "test3"),
                             names_to = "test", 
                             values_to = "ODR") |>
  left_join(tidy_opt_co) |>
  mutate(neg = as.numeric(ODR < co))

N_per_species <- group_by(kiteval, species) |>
  summarise(N = n())
```
