## Supplementary material for "Assessing measurement uncertainty at laboratory network scale and its impact on diagnostic performance in the absence of a gold standard: application to ELISA tests for *Coxiella burnetii in ruminants*": code and data: README.txt

Assessing measurement uncertainty at laboratory network scale and its impact on diagnostic performance in the absence of a gold standard: 
application to ELISA tests for Coxiella burnetii in ruminants

L. Rivière, M.L. Delignette Muller, M. Prigent, T. Lurier, E. Rousset
_______________________________________________________________________________________________________________________________________

To reproduce all the analyses and figures, run the files in this order:
1. outlier_detection.R
2. measurement_uncertainty_assessment.R
3. changing_result_probability.R
4. PPP_calculation.R
5. Se_Sp_estimation.R

Intermediate results are provided so that you can run the scripts independently. They are stored in the "intermediate_results" subfolder.
Figures and tables will be saved in the "results" subfolder.

The "data" subfolder contains the data sets, scripts to load them, and scripts with functions needed in the main scripts. 
You don't need to run these scripts, as they are loaded in the main code files via the function source().

The ILPT.csv file contains the ILPT data. The data are describe in Rousset et al. 2023: https://anses.hal.science/anses-05126539
The kiteval4500.csv file contains simulated data similar to the realKiteval4500 data. 
Therefore, the results obtained by runing these code files will be slightly different from those shown in the article.
_______________________________________________________________________________________________________________________________________

The code was run in R with the following software versions:

R version 4.2.2 (2022-10-31 ucrt)
Platform: x86_64-w64-mingw32/x64 (64-bit)
Running under: Windows 10 x64 (build 19045)

Matrix products: default

locale:
[1] LC_COLLATE=French_France.utf8  LC_CTYPE=French_France.utf8    LC_MONETARY=French_France.utf8 LC_NUMERIC=C                  
[5] LC_TIME=French_France.utf8    

attached base packages:
[1] parallel  stats     graphics  grDevices utils     datasets  methods   base     

other attached packages:
 [1] ggtext_0.1.2      gridExtra_2.3     cubature_2.1.0    lattice_0.20-45   runjags_2.2.2-1.1 rjags_4-15        coda_0.19-4      
 [8] ggplot2_3.5.1     dplyr_1.1.4       tidyr_1.3.1       lme4_1.1-35.5     Matrix_1.5-1     

loaded via a namespace (and not attached):
 [1] Rcpp_1.0.10      pillar_1.9.0     compiler_4.2.2   nloptr_2.1.1     tools_4.2.2      boot_1.3-28      lifecycle_1.0.3 
 [8] tibble_3.2.1     nlme_3.1-160     gtable_0.3.1     pkgconfig_2.0.3  rlang_1.1.3      cli_3.6.0        rstudioapi_0.14 
[15] xml2_1.3.6       withr_3.0.2      generics_0.1.3   vctrs_0.6.5      gridtext_0.1.5   grid_4.2.2       tidyselect_1.2.1
[22] glue_1.6.2       R6_2.5.1         fansi_1.0.4      minqa_1.2.8      purrr_1.0.1      magrittr_2.0.3   scales_1.3.0    
[29] MASS_7.3-58.1    splines_4.2.2    colorspace_2.1-0 utf8_1.2.2       munsell_0.5.0 
