## Supplementary figures and images for "Assessing measurement uncertainty at laboratory network scale and its impact on diagnostic performance in the absence of a gold standard: application to ELISA tests for *Coxiella burnetii in ruminants*"

### fig3.jpg

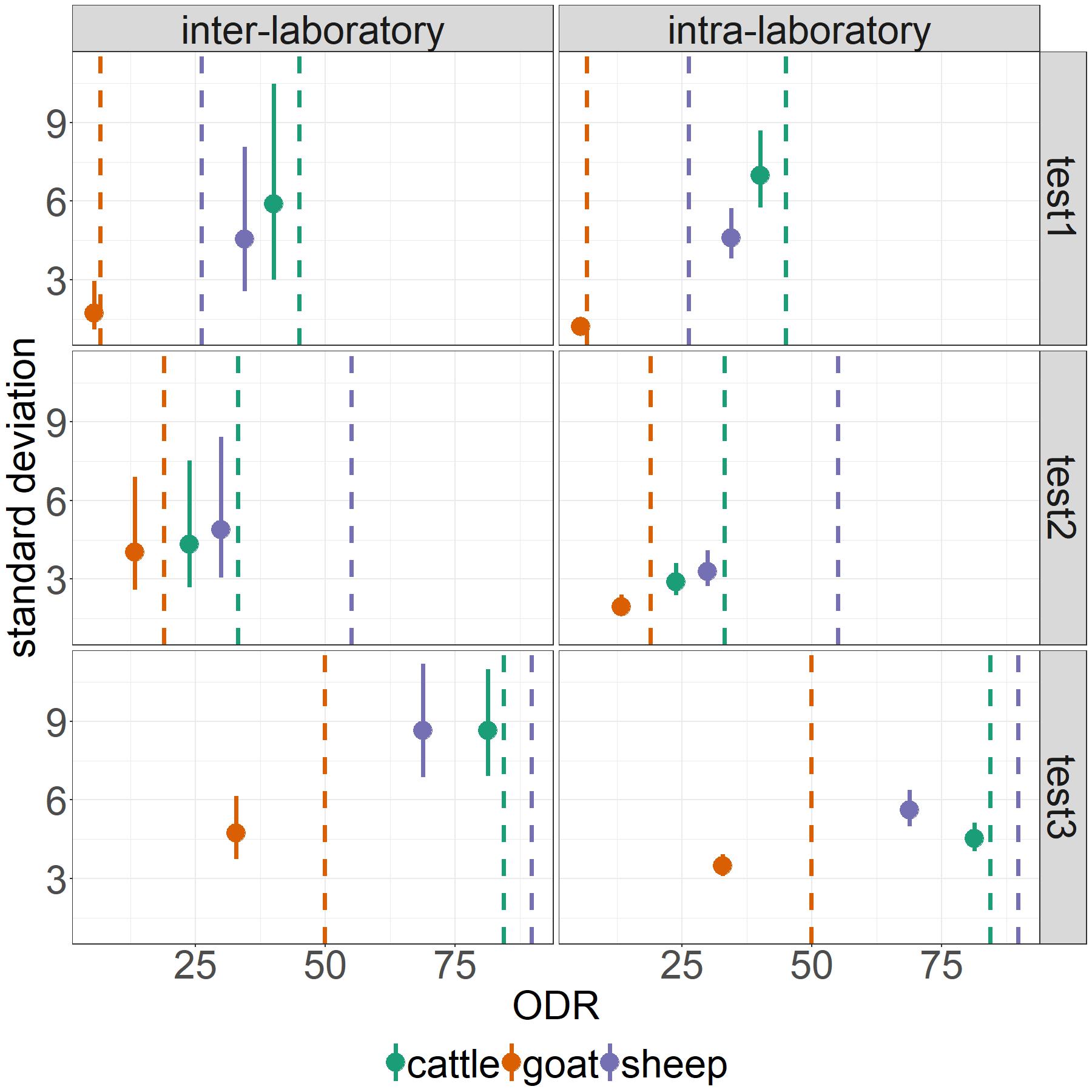

### fig4.jpg

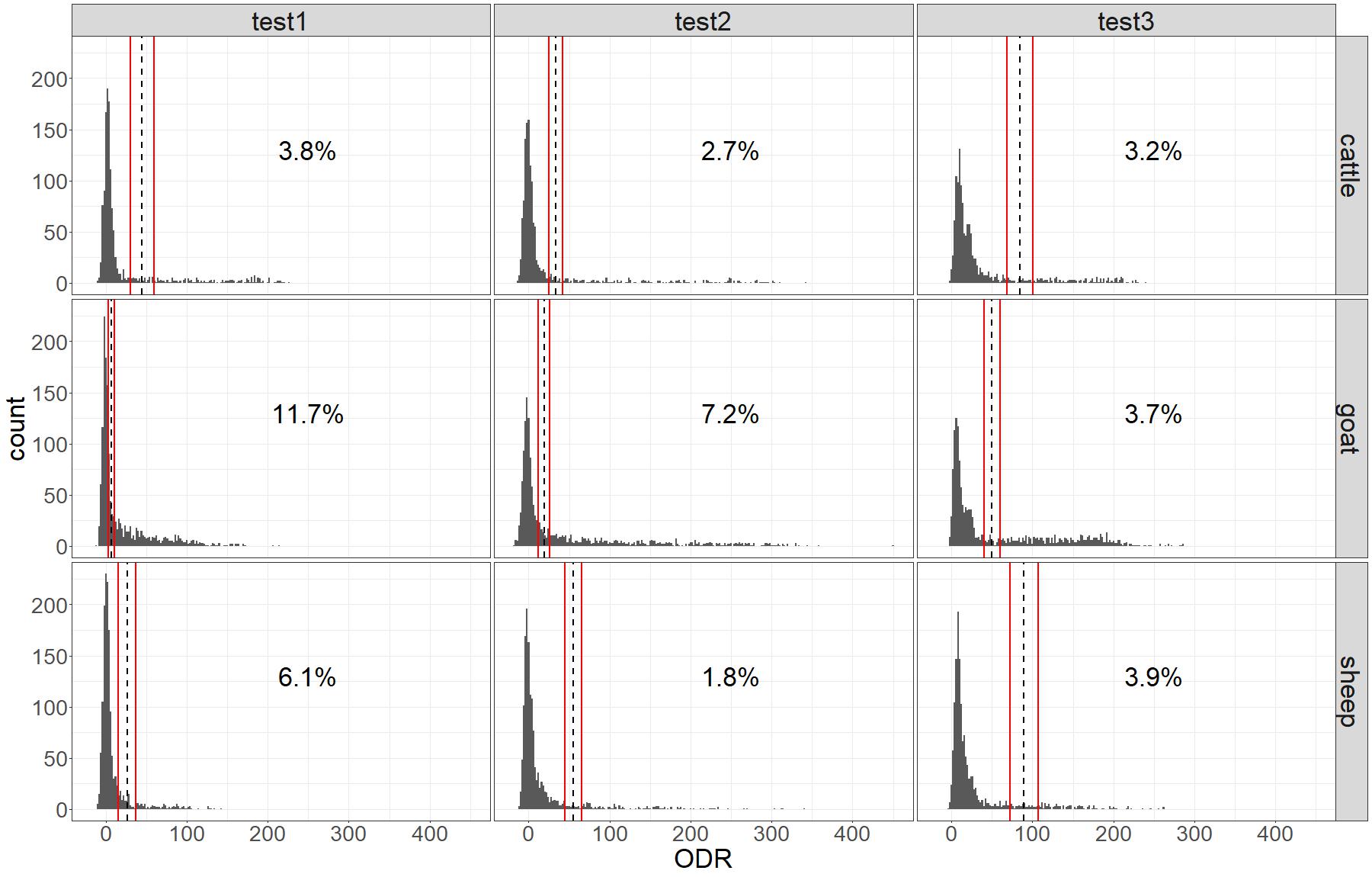

### fig6.jpg

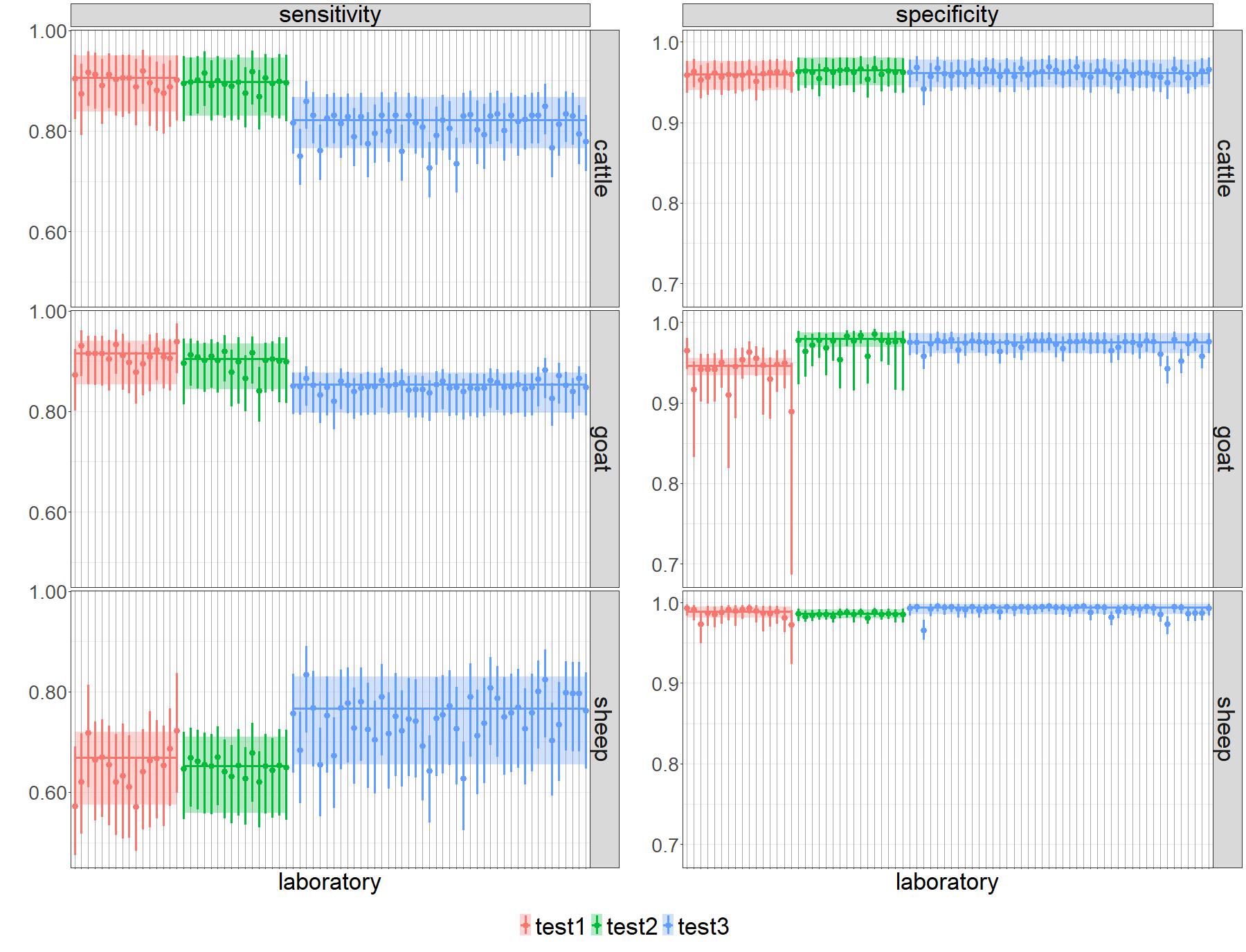

### fig7.jpg

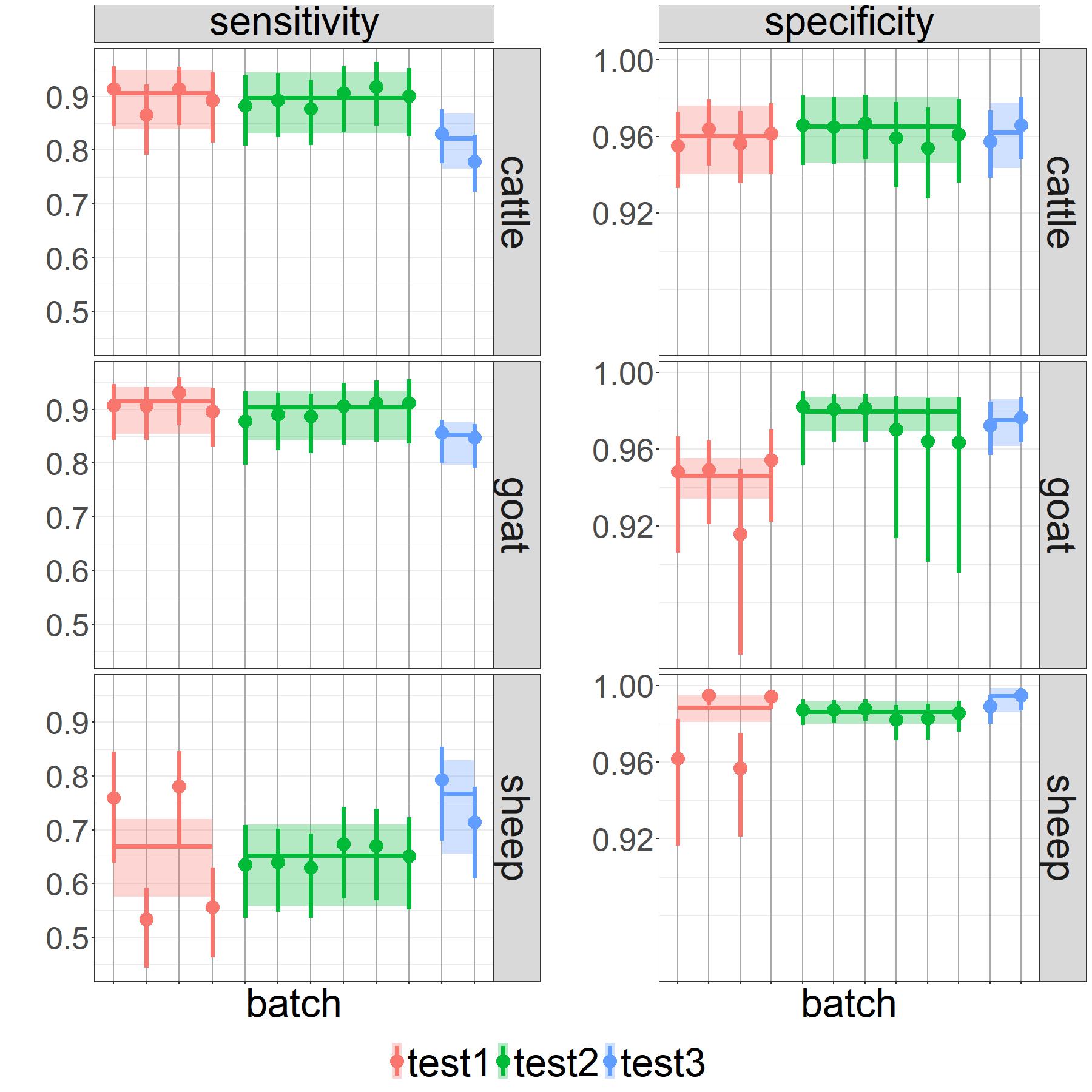

### figA1.2.jpg

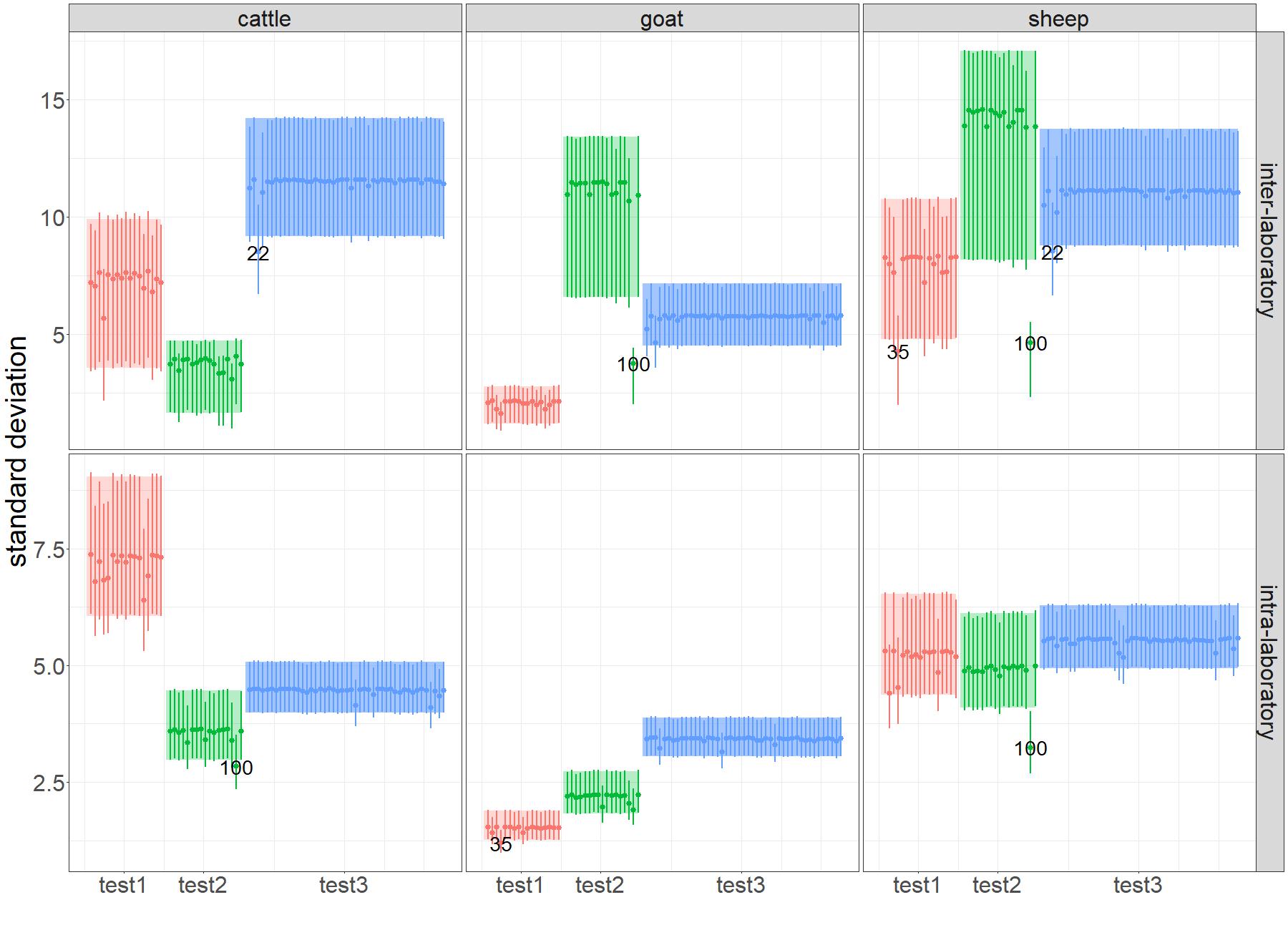

### figA5.jpg

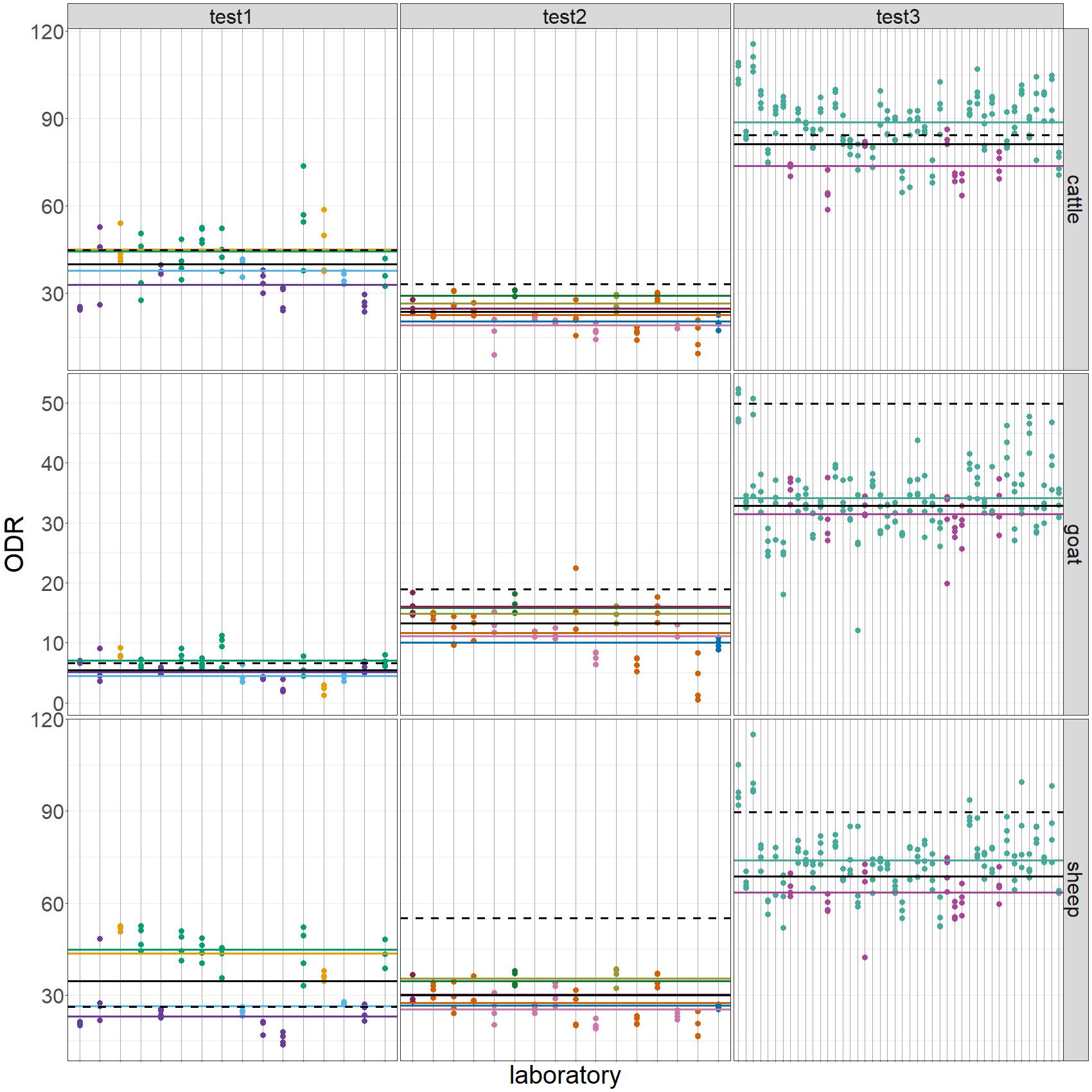
