## appendix for "Assessing measurement uncertainty at laboratory network scale and its impact on diagnostic performance in the absence of a gold standard: application to ELISA tests for *Coxiella burnetii in ruminants*"

### A1. Detection of outlier laboratories

The method used to detect outlier laboratories is illustrated in Figure A1.1. It was applied to each test-serum combination.

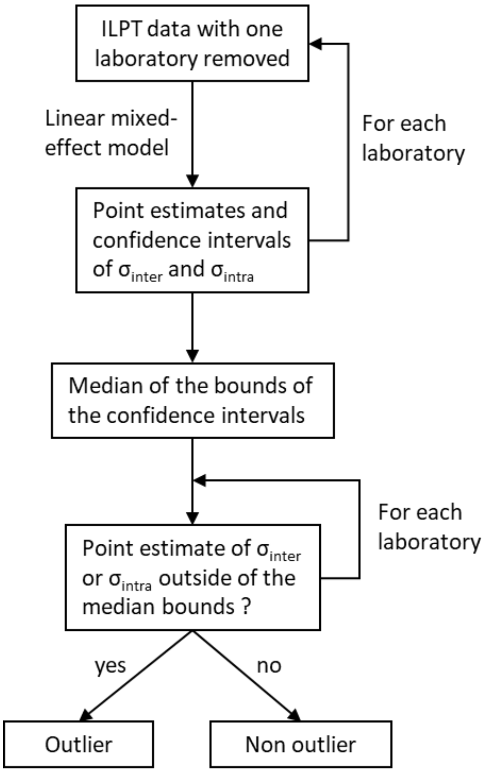


Figure A1.1: Method used for the detection of outlier laboratories

The results of the outlier detection process are illustrated in Figure A1.2. Three laboratories (one per test) were identified as outliers.

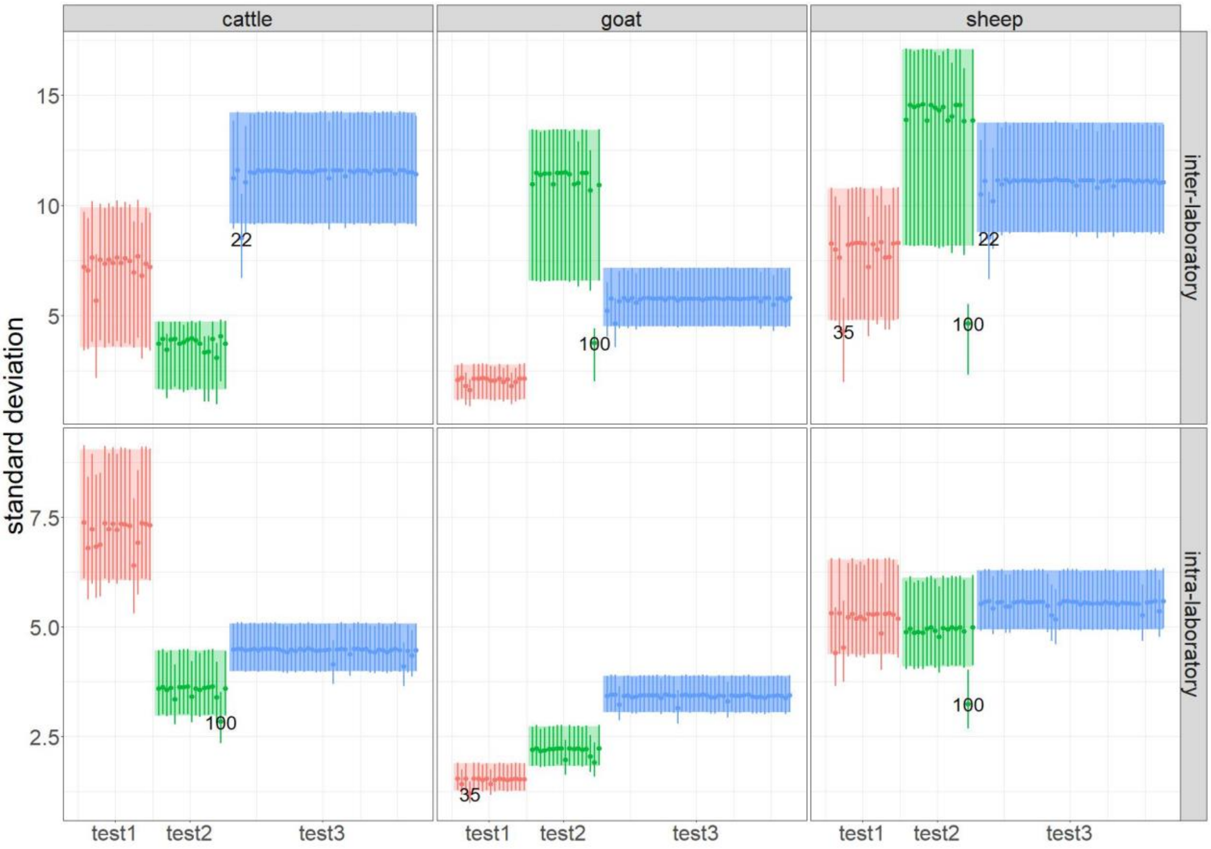


Figure A1.2: Results from the outlier detection process. The points and vertical lines represent the point estimates and 95% confidence intervals of the inter- and intra-laboratory standard deviations. The coloured bands represent the median confidence intervals. The numbers represent the identification number of the laboratories for which, when removed, the point estimate is outside of the median bounds of the confidence intervals.

### A2. Prior choice

Gelman (2006) recommends using a half-Cauchy distribution centred on 0 as the prior for standard deviations describing random effects in multilevel models, with a scale parameter corresponding to a realistic upper bound for the standard deviation. Based on the previous ILPTs for Q fever ELISA tests (such as Rousset & Dufour, 2019), the inter-laboratory standard deviation was expected to be below 20. Therefore, a half-Cauchy distribution with a scale parameter of 20 was used as a prior for the inter-laboratory standard deviations: $\sigma_{inter} \sim \text{half-Cauchy}(0{,}20)$. The intra-laboratory standard deviation was not assessed in the previous ILPTs, but was expected to be less or equal to the inter-laboratory standard deviation. Therefore, the same prior was used on the intra-laboratory standard deviation: $\sigma_{intra} \sim \text{half-Cauchy}(0{,}20)$.

Broad Gaussian distributions were chosen as priors for the mean ODR and the batch effects, in order to avoid restricting their estimates too much. The prior for the serum mean ODR was centred on the mean ODR of the corresponding serum measured by the national Q fever laboratory $\mu$ (Rousset & Prigent, 2023), with a standard deviation of the same order of magnitude as the expected upper bound for inter- and intra-laboratory standard deviation: $x_0 \sim \text{Normal}(\mu, 20)$.

The prior for the batch effects was centred on 0. The data from the monitoring of the batches between 2012 and 2022 (Rousset *et al.*, 2023) were used to choose the standard deviation of this prior. For this monitoring, several samples of two reference sera were analysed with each new marketed batch. The maximum of the empirical standard deviation across all tests and reference sera, multiplied by 2, was chosen as the prior standard deviation: $b_k \sim \text{Normal}(0, 12.4)$

### A3. PPP calculation

Let $S^+$(respectively $S^-$) represent the event of being truly seropositive (respectively seronegative), and let $P_t$ be the random variable equal to 1 if test $t$ is positive and 0 if it is negative.

For T tests, the probability of an individual being truly seropositive given its test results is:

$$PPP = \text{P}(S^+|P_1 = p_1, \dots, P_T = p_T)$$

$$= \frac{\text{P}(S^+) \times \text{P}(P_1 = p_1, \dots, P_T = p_T|S^+)}{\text{P}(P_1 = p_1, \dots, P_T = p_T)}$$

$$= \frac{\text{P}(S^+) \times \text{P}(P_1 = p_1, \dots, P_T = p_T|S^+)}{\text{P}(S^+) \times \text{P}(P_1 = p_1, \dots, P_T = p_T|S^+) + \text{P}(S^-) \times \text{P}(P_1 = p_1, \dots, P_T = p_T|S^-)}$$

In the case of conditionally dependent tests,

$$\text{P}(P_1 = p_1, \dots, P_T = p_T|S^+) = \prod_{t=1}^{T} \text{P}(P_t = p_t|S^+) + \gamma_{Se\ p_1,\dots,p_T}$$

and $\text{P}(P_1 = p_1, \dots, P_T = p_T|S^-) = \prod_{t=1}^{T} \text{P}(P_t = p_t|S^-) + \gamma_{Sp\ p_1,\dots,p_T}$

Where $\gamma_{Se\ p_1,\dots,p_T}$ (respectively $\gamma_{Sp\ p_1,\dots,p_T}$) represents the excess of probability, compared to the expected probability under the assumption of conditional independence between tests, for a truly seropositive (respectively seronegative) individual to have the results $p_1, \dots, p_T$.

For each test $t$, $\text{P}(P_t = 1|S^+) = Se_t$ and $\text{P}(P_t = 0|S^+) = (1 - Sp_t)$, where $Se_t$ and $Sp_t$ are the sensitivity and specificity of test $t$.

Therefore, $\text{P}(P_1 = p_1, \dots, P_T = p_T|S^+) = \prod_{t=1}^{T}(p_t \times Se_t + (1 - p_t) \times (1 - Se_t)) + \gamma_{Se\ p_1,\dots,p_T}$

Similarly, $\text{P}(P_1 = p_1, \dots, P_T = p_T|S^-) = \prod_{t=1}^{T}(p_t \times (1 - Sp_t) + (1 - p_t) \times Sp_t) + \gamma_{Sp\ p_1,\dots,p_T}$

Therefore, we calculated the PPPs in each herd for each test result combination using the following formulas:

$$PPP_{h,000} = \frac{prev_h \times ((1 - Se_1) \times (1 - Se_2) \times (1 - Se_3) + \gamma_{Se\ 000})}{prev_h \times \left((1 - Se_1) \times (1 - Se_2) \times (1 - Se_3) + \gamma_{Se\ 000}\right) + (1 - prev_h) \times (Sp_1 \times Sp_2 \times Sp_3 + \gamma_{Sp\ 000})}$$

$$PPP_{h,001} = \frac{prev_h \times ((1 - Se_1) \times (1 - Se_2) \times Se_3 + \gamma_{Se\ 001})}{prev_h \times \left((1 - Se_1) \times (1 - Se_2) \times Se_3 + \gamma_{Se\ 001}\right) + (1 - prev_h) \times (Sp_1 \times Sp_2 \times (1 - Sp_3) + \gamma_{Sp\ 001})}$$

$$PPP_{h,010} = \frac{prev_h \times ((1 - Se_1) \times Se_2 \times (1 - Se_3) + \gamma_{Se\ 010})}{prev_h \times \left((1 - Se_1) \times Se_2 \times (1 - Se_3) + \gamma_{Se\ 010}\right) + (1 - prev_h) \times (Sp_1 \times (1 - Sp_2) \times Sp_3 + \gamma_{Sp\ 010})}$$

$$PPP_{h,011} = \frac{prev_h \times ((1 - Se_1) \times Se_2 \times Se_3 + \gamma_{Se\ 011})}{prev_h \times \left((1 - Se_1) \times Se_2 \times Se_3 + \gamma_{Se\ 011}\right) + (1 - prev_h) \times (Sp_1 \times (1 - Sp_2) \times (1 - Sp_3) + \gamma_{Sp\ 011})}$$

$$PPP_{h,100} = \frac{prev_h \times (Se_1 \times (1 - Se_2) \times (1 - Se_3) + \gamma_{Se\ 100})}{prev_h \times (Se_1 \times (1 - Se_2) \times (1 - Se_3) + \gamma_{Se\ 100}) + (1 - prev_h) \times ((1 - Sp_1) \times Sp_2 \times Sp_3 + \gamma_{Sp\ 100})}$$

$$PPP_{h,101} = \frac{prev_h \times (Se_1 \times (1 - Se_2) \times Se_3 + \gamma_{Se\ 101})}{prev_h \times (Se_1 \times (1 - Se_2) \times Se_3 + \gamma_{Se\ 101}) + (1 - prev_h) \times ((1 - Sp_1) \times Sp_2 \times (1 - Sp_3) + \gamma_{Sp\ 101})}$$

$$PPP_{h,110} = \frac{prev_h \times (Se_1 \times Se_2 \times (1 - Se_3) + \gamma_{Se\ 110})}{prev_h \times (Se_1 \times Se_2 \times (1 - Se_3) + \gamma_{Se\ 110}) + (1 - prev_h) \times ((1 - Sp_1) \times (1 - Sp_2) \times Sp_3 + \gamma_{Sp\ 110})}$$

$$PPP_{h,111} = \frac{prev_h \times (Se_1 \times Se_2 \times Se_3) + \gamma_{Se\ 111})}{prev_h \times (Se_1 \times Se_2 \times Se_3 + \gamma_{Se\ 111}) + (1 - prev_h) \times ((1 - Sp_1) \times (1 - Sp_2) \times (1 - Sp_3) + \gamma_{Sp\ 111})}$$

Where $PPP_{h,abc}$ is the posterior positive probability for an individual in herd $h$ whose results for tests 1, 2 and 3 are $a$, $b$ and $c$ respectively (0 represents a negative result and 1 a positive result); $prev_h$ is the prevalence in herd $h$; $Se_1$, $Se_2$, $Se_3$, $Sp_1$, $Sp_2$ and $Sp_3$ are the estimated sensitivities and specificities of tests 1, 2 and 3 respectively; and $\gamma_{Se\ abc}$ and $\gamma_{Sp\ abc}$ are the conditional dependence terms.

### A4. Number of false negatives and false positives with and without measurement uncertainty

Table A4: Number of false negatives and false positives taking and not taking into account measurement uncertainty per 1000 individuals (point estimate and 95% credibility interval)

| Species | Test | False negatives | | False positives | |
|---|---|---|---|---|---|
| | | without measurement uncertainty | with measurement uncertainty | without measurement uncertainty | with measurement uncertainty |
| Cattle | Test 1 | 13.7 [9.6; 21.9] | 15.0 [10.9; 23.1] | 27.5 [21.2; 34.9] | 29.5 [23.3; 37.0] |
| | Test 2 | 11.6 [7.3; 19.7] | 13.2 [9.0; 21.2] | 22.2 [15.8; 29.7] | 24.5 [18.0; 32.0] |
| | Test 3 | 21.0 [17.7; 28.6] | 23.5 [20.2; 31.2] | 23.6 [17.0; 31.1] | 25.4 [19.0; 32.6] |
| Goat | Test 1 | 38.8 [29.1; 68.7] | 38.8 [29.7; 67.3] | 7.9 [5.1; 12.7] | 15.1 [10.8; 25.4] |
| | Test 2 | 41.9 [31.9; 71.7] | 46.2 [36.4; 74.8] | 7.8 [5.1; 12.0] | 12.2 [8.5; 19.0] |
| | Test 3 | 69.2 [60.6; 98.5] | 69.5 [61.3; 98.2] | 15.4 [11.0; 21.2] | 18.0 [13.5; 23.8] |
| Sheep | Test 1 | 44.7 [37.7; 71.2] | 45.6 [38.7; 71.5] | 6.1 [3.0; 10.8] | 11.9 [7.8; 19.3] |
| | Test 2 | 44.9 [37.0; 71.5] | 45.2 [37.3; 71.9] | 10.6 [6.5; 14.1] | 10.8 [7.0; 14.2] |
| | Test 3 | 30.4 [22.5; 56.2] | 33.1 [25.3; 58.7] | 2.0 [0.4; 6.5] | 5.0 [3.3; 9.2] |

### A5. ILPT data

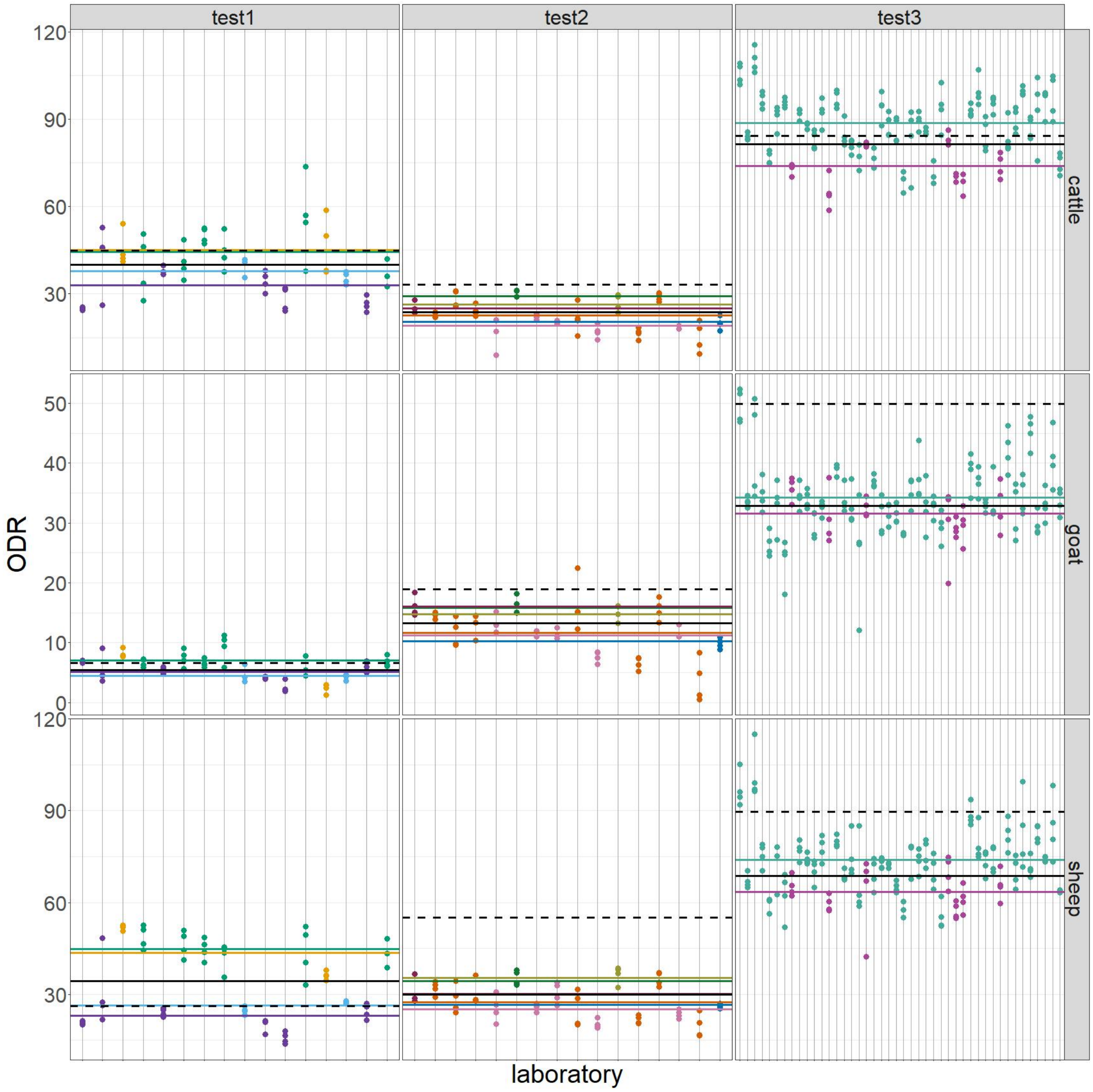


Figure A5: ILPT data and estimations from the linear mixed-effects model. Each point represents the observed ODR of a sample. The black solid lines represent the serum mean ODR ($x_0$). The coloured lines represent the batch ODR ($x_0 + b_k$), where each colour represents a batch. The dashed lines represent the cut-off.
